# Wireless closed-loop smart bandage for chronic wound management and accelerated tissue regeneration

**DOI:** 10.1101/2022.01.16.476432

**Authors:** Yuanwen Jiang, Artem A. Trotsyuk, Simiao Niu, Dominic Henn, Kellen Chen, Chien-Chung Shih, Madelyn R. Larson, Alana M. Mermin-Bunnell, Smiti Mittal, Jian-Cheng Lai, Aref Saberi, Ethan Beard, Serena Jing, Donglai Zhong, Sydney R. Steele, Kefan Sun, Tanish Jain, Eric Zhao, Christopher R. Neimeth, Willian G. Viana, Jing Tang, Dharshan Sivaraj, Jagannath Padmanabhan, Melanie Rodrigues, David P. Perrault, Arhana Chattopadhyay, Zeshaan N. Maan, Melissa C. Leeolou, Clark A. Bonham, Sun Hyung Kwon, Hudson C. Kussie, Katharina S. Fischer, Gurupranav Gurusankar, Kui Liang, Kailiang Zhang, Ronjon Nag, Michael P. Snyder, Michael Januszyk, Geoffrey C. Gurtner, Zhenan Bao

## Abstract

Chronic non-healing wounds represent a major source of morbidity for patients and a significant economic burden. Current wound care treatments are generally passive and are unable to adapt to changes in the wound environment in real time. By integrating multimodal sensors and adding stimulators in a bandage, real-time physiological monitoring is possible and provides an opportunity for active intervention into the complex wound environment. Here, we develop a battery-free flexible bioelectronic system consisting of wirelessly powered, closed-loop sensing and stimulation circuits with tissue-interfacing tough conducting hydrogel electrodes for robust signal transduction, on-demand adhesion, and detachment. Using multiple pre-clinical models, we demonstrate the capability of our wound care system to continuously monitor skin impedance and temperature, to trigger directional electrical stimulation. The accelerated wound closure was confirmed to be due to the activation of pro-regenerative genes linked to accelerated wound closure, increased neovascularization, and enhanced dermal recovery.

## Introduction

Chronic non-healing wounds represent a significant healthcare burden, with more than 6 million individuals affected in the United States alone^1^. These wounds are associated with loss of function and mobility, increased social stress and isolation, depression and anxiety, prolonged hospitalization, and overall increased morbidity and mortality. In addition, the financial cost to the healthcare system for the management of chronic wound-related complications has been estimated to exceed $25 billion annually^1^. A chronic wound is defined as a wound that has failed to heal by 8-12 weeks and is unable to restore function and anatomical integrity to the affected site^2^.

In normal wound healing, when an injury occurs, the tissue undergoes three canonical stages of wound regeneration: inflammation, new tissue formation, and remodeling^3^. During each stage, different cells are recruited, migrate, become activated, and proliferate to achieve tissue regeneration and reduce infection^4^. When this carefully orchestrated process is impaired, there is often not typically a single cause, but rather multiple factors, that contribute. These factors include comorbidities such as diabetes, infection, ischemia, metabolic conditions, immunosuppression, and radiation, which can result in high level of proteases, elevated inflammatory markers, low growth factor activity, and reduced cellular proliferation within the wound bed. This can lead to significant patient discomfort and increased hospitalization rates^5,6^.

While interventions for chronic wounds exist, such as growth factors, extracellular matrix, engineered skin, and negative pressure wound therapy, these treatments are only moderately effective^6,7^. Current standard-of-care wound dressings are passive and do not actively respond to variations in the wound environment. Smart bandage technologies are well positioned to address these challenges with their ability to integrate multimodal sensors and stimulators for real-time monitoring and active wound care treatment with minimal physician intervention^8,9^.

Prior research has demonstrated that as a wound heals, skin impedance increases^10^. When a wound becomes infected, however, wound impedance sharply oscillates due to the development of biofilm^11^. As the infection develops further, local inflammation increases wound temperature^12^. Both signals can be easily captured by low-cost sensors embedded in a wearable device to act as a sentinel for impending wound infection. These biophysical signals provide rapid, robust, and accurate information about wound conditions in real time, creating an opportunity to diagnose and monitor a non-healing wound quickly and autonomously in a closed-loop fashion.

Current smart bandage technologies have demonstrated promise in their ability to sense physiological conditions. This includes detecting pH^13^, temperature^14,15^, oxygenation^16^, impedance^10,17^, motions^18^, and enzymatic fluctuations^19^ of the wound. It has also been well established that electrical stimulation can reduce bacterial colonization, biofilm infection and restore normal wound healing *in vivo*^20^. Moreover, electrical stimulation has also been shown to improve tissue perfusion, stimulate immune cell function, and accelerate keratinocyte migration through a process known as galvanotaxis^21,22^. Unfortunately, current electrical stimulation devices are bulky, tethered by wires, and uncomfortable to wear, limiting patient compliance. In addition, there have not been significant advancements in incorporating both sensing and electrical stimulation technologies to simultaneously deliver active wound care (**Table S1**). There remains a need to develop portable, autonomous, inexpensive devices to improve wound care.

For improved therapeutic outcomes, an ideal smart bandage platform needs to meet the following requirements. First, it needs to be flexible and wirelessly operated to avoid any undesired tethering and discomfort caused by conventional rigid, battery powered devices. Next, it should integrate both sensing and stimulation modalities for autonomous, closed-loop wound management. Finally, it should have on-demand skin adhesion with a tight interface for robust signal transduction and energy delivery during operation, while providing easy detachment to avoid possible secondary skin damage during device removal.

To address these requirements, we developed a battery-free flexible bioelectronic system consisting of wirelessly powered sensing and stimulation circuits with tissue-interfacing tough hydrogel electrodes using a biocompatible conducting polymer. We anticipate that this smart bandage will improve therapeutic outcomes and provide new knowledge for wound care.

Specifically, we designed a miniaturized flexible printed circuit board (FPCB) containing an energy harvesting antenna, a microcontroller unit, a crystal oscillator, and filter circuits for dual channel continuous sensing of wound impedance and temperature, as well as a parallel stimulation circuit to deliver programmed electrical cues for accelerated wound healing. To ensure efficient signal exchange and energy delivery between the circuits and the soft skin tissue, we designed a low-impedance and adhesive hydrogel electrode based on poly(3,4-ethylenedioxythiophene):polystyrene sulfonate (PEDOT:PSS). Compared to well-established ionically conducting hydrogels, our dual-conducting (i.e. both electrically and ionically conductive) hydrogel has lower impedance across the entire frequency domain, giving rise to more efficient charge injection during stimulation^23,24^. To mitigate secondary skin damage when peeling off the adhesive electrodes, we introduced a thermally controlled reversible phase transition mechanism to the hydrogel backbone and achieved two orders of magnitude lower adhesion at elevated temperature when compared to the normal skin temperature. Using multiple pre-clinical wound and disease models, we found that our wound care system could deliver directional electrical cues, leading to the activation of pro-regenerative genes linked to accelerated wound closure, increased neovascularization, and enhanced dermal recovery.

### System overview

Our integrated wound management system consists of a battery-free, wirelessly powered FPCB for simultaneous wound treatment and monitoring, as well as a tissue-interfacing conducting adhesive hydrogel interface for robust and gentle skin integration (**Fig. 1a-b**). Due to the thin layout of the FPCB (~100 μm board thickness) and low modulus of the gel interface, the smart bandage is flexible and can be conformably attached to wound surfaces (**Fig. 1c-e**). With an antenna coil that resonates at 13.56 MHz, our smart bandage can be inductively coupled with an external radiofrequency identification (RFID) reader. Through the RF energy harvesting process, the antenna can provide power to apply electric bias across the wound for programmed treatment and, at the same time, drive the microcontroller unit (MCU) and other integrated circuits (e.g., oscillator and filter), for continuous monitoring of wound impedance and temperature via a near-field communication (NFC) transponder in the MCU under the ISO15693 protocol (**Fig. 1f, 2a**).

**Figure 1.**
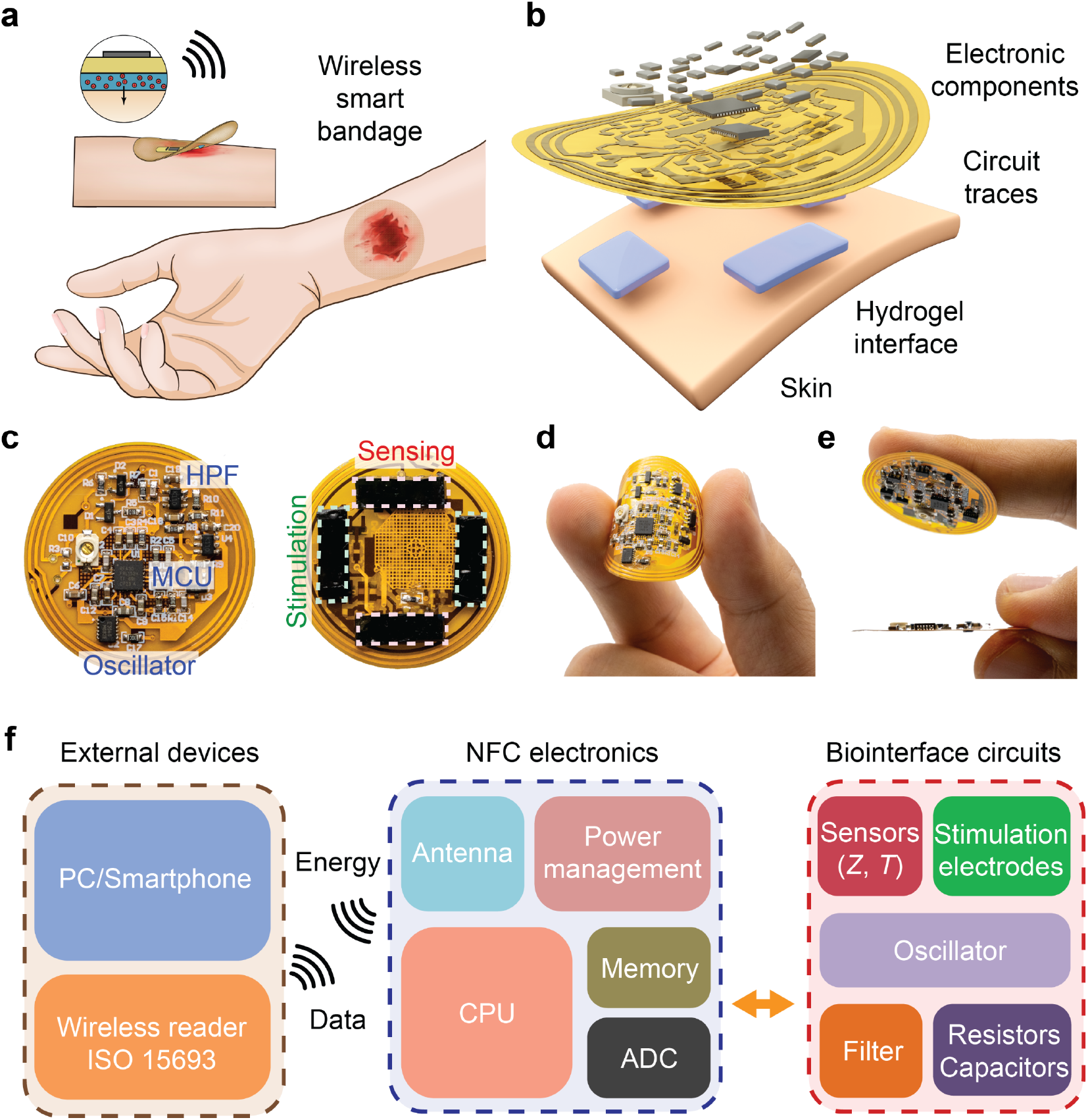
Overall design of the wireless smart bandage for chronic wound management. **a** and **b,** Schematic diagram (**a**) and exploded view (**b**) of the wireless smart bandage including flexible printed circuit board (FPCB) and tissue-interfacing conducting adhesive hydrogel. **c**, Photographs of the front (left) and back (right) sides of the smart bandage showing the microcontroller unit (MCU), crystal oscillator, high-pass filter (HPF), and stimulation and sensing electrodes. **d** and **e**, Photographs showing the flexibility of the FPCB (**d**), adhesion of the hydrogel interface, and the thin layout of the board (**e**, lower). **f**, Block diagram illustrating the key components of the wireless smart bandage system composed of near-field communication (NFC) electronics with parallel stimulation and sensing modalities. CPU, central processing unit; ADC, analog-digital converter.

**Figure 2.**
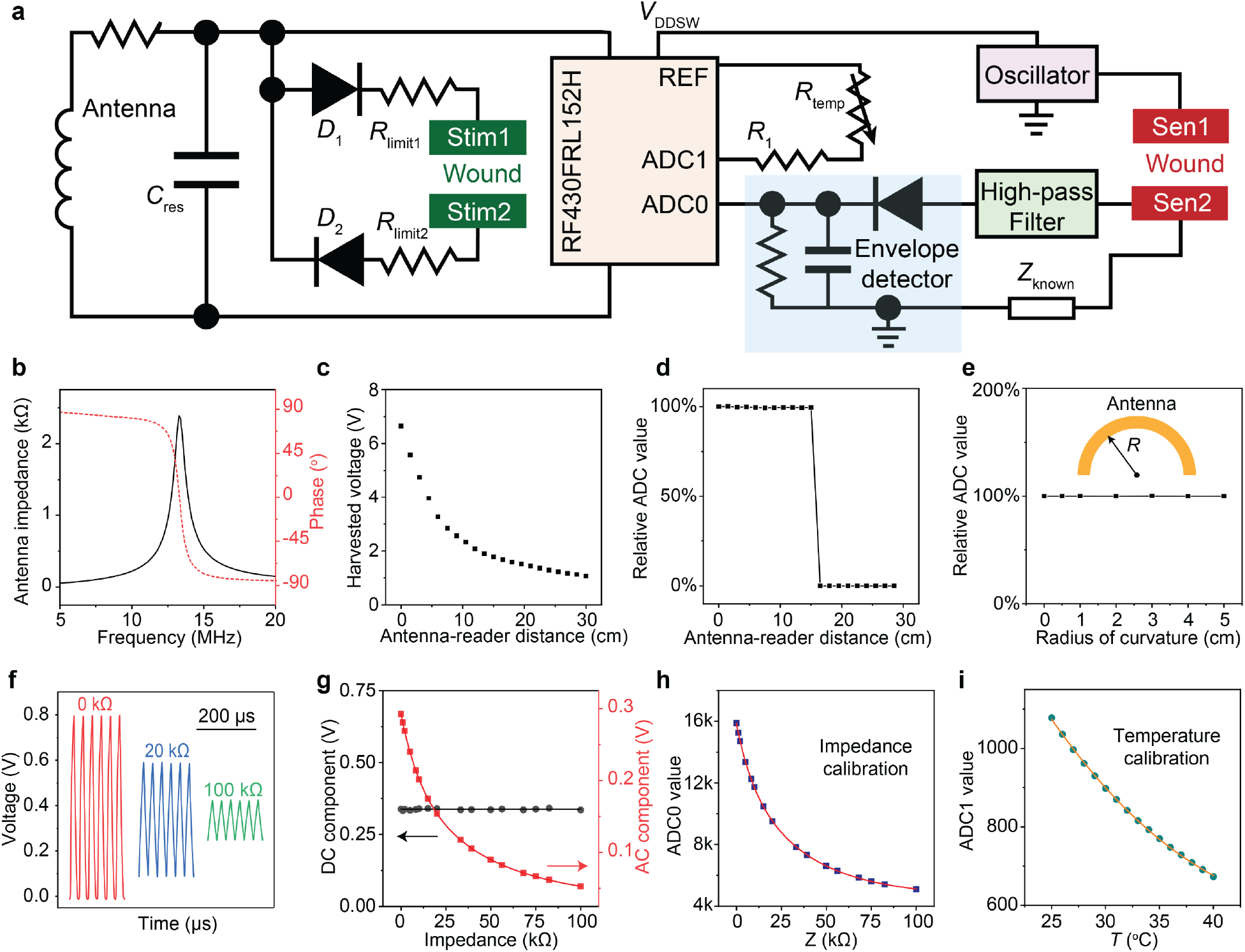
Validation of the wireless sensing and stimulation circuits. **a**, Circuit diagram of the wireless smart bandage for simultaneous sensing and stimulation. **b**, Antenna resonant frequency and quality factor as measured by a vector network analyzer (VNA). **c**, Measured RF harvested voltage as a function of antenna-reader distance. **d**, Wireless readout operation from the microcontroller can function stably up to 15 cm away from the external reader. **e**, Wireless sensing can remain stable with bending radius down to 0.5 cm. **f** and **g**, Voltage output after the high-pass filter showing reduced AC amplitudes with respect to larger resistance values. In the meantime, the DC component of the signals remain constant for all resistors tested. **h** and **i**, Calibration curves of ADC values under known impedance (**h**) and temperature (**i**).

### Wireless circuit design

For the wireless antenna, we designed a 5-turn coil with an optimum inductance of ~1.5 μH, offering a high RF harvested voltage and wide tunability to reach a resonant frequency of 13.56 MHz for maximized wireless communication signal gain (**Fig. 2a-c, Fig. S1**). Additionally, the quality factor (*Q*) of the antenna is ~18, which strikes the balance between energy harvesting efficiency and wireless communication bandwidth (**Fig. 2b**). As a result, our antenna offers a wide and stable 15 cm wireless communication distance (**Fig. 2c-d**). Our device function also remained stable upon bending (**Fig. 2e, Fig. S3**).

The NFC transponder we used (RF430FRL152H), offers two 14-bits analog-digital converters (ADCs) to serve as the analog front-end interface. To best monitor the condition of the wound, we chose to integrate two sensors (one thermistor and one impedance sensor), which serve as good proxies for determining infection and inflammatory states of the wound^10,17,25^. The RF430FRL152H transponder has a direct thermistor support (ADC1 channel) by emitting a small μA level current on the thermistor and sampling the voltage. For impedance sensing, an oscillator was used to generate a 32.768 kHz square wave alternating current (AC) signal (**Fig. S2**) that passed through the wound, and a known impedance component (*Z*_known_). Through a voltage divider, the AC signal applied on *Z*_known_ could then reflect the wound impedance **(Fig. 2f, Fig. S2)**. This received AC signal was further conditioned through a high-pass filter to remove the random direct current (DC) component inside the oscillation signal (**Fig. 2g, Fig. S2**). Finally, an envelope detector was used to convert the AC signal amplitude to a DC voltage, which was captured by the ADC0 channel inside the RF430FRL152H transponder. With standard impedance components and controlled temperature, we calibrated both ADC channels in our integrated design (**Fig. 2h, i**). Moreover, due to the use of different frequency bands and proper signal conditioning, the stimulation channel had no interference with the sensing channel (**Fig. S4**). Finally, the sensing data could be analyzed in real-time to provide feedback on the stimulation pattern for closed-loop operation (**Fig. S5**).

### Hydrogel interface

To ensure an intimate skin interface and robust electrical communication between the circuit and tissue through the soft hydrogel, the gel electrode interface should have the following characteristics: low contact impedance, high toughness, and tunable adhesion. The low contact impedance is to ensure sensitive sensing and efficient charge injection by electrical stimulation.

The high toughness requirement is to avoid mechanical damage during motion. Finally, the tissue interfacing gel needs to have on-demand adhesion to the wound tissue to provide good adhesion during therapy while also easy, gentle removal upon external triggers (e.g., gentle heating) to mitigate secondary damage to the delicate wounded tissue and prevent a commonly occurred skin condition known as medical adhesive-related skin injury^26,27^ (**Fig. 3a**).

**Figure 3.**
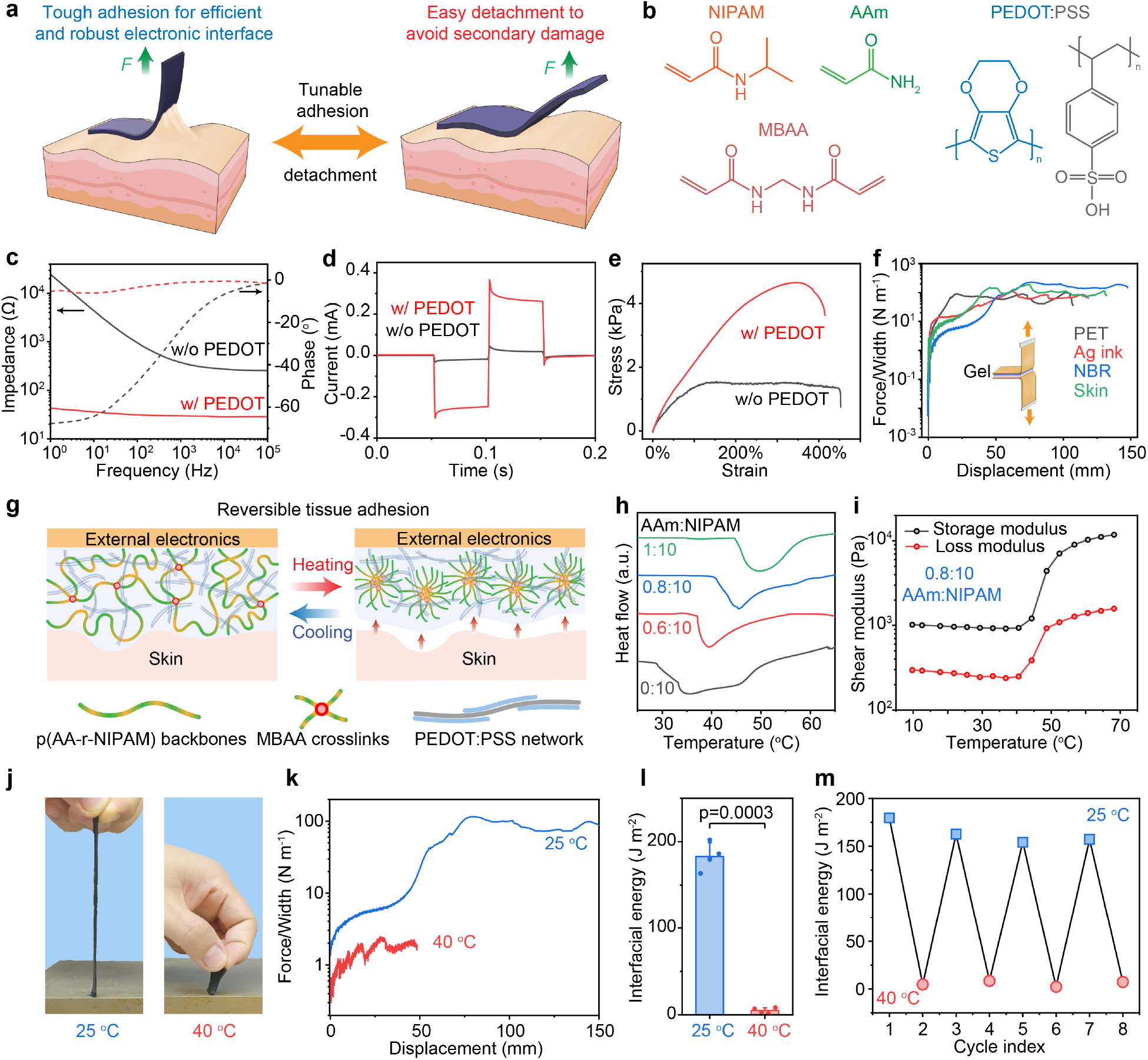
Tough and low-impedance conductive hydrogel electrode with reversible tissue adhesion. **a,** Schematic diagram illustrating the requirements for the hydrogel interface in the smart bandage. During device operation, the hydrogel electrode needs to possess simultaneously high toughness and adhesion to avoid damage or detachment. When peeling off the device after the treatment period, the tissue interfacing gel needs to be easily detachable to minimize secondary damage to the delicate wounded tissue. **b**, Molecular structures of the monomers, crosslinker, and conducting polymer for the interpenetrated double network. **c** and **d**, Electrochemical impedance spectroscopy (EIS, **c**) and chronoamperometry (**d**) of hydrogels with (20 mg mL^-1^ PEDOT:PSS with 150 mg mL^-1^ NIPAM and 12 mg mL^-1^ AAm) and without (150 mg mL^-1^ NIPAM and 12 mg mL^-1^ AAm only) PEDOT:PSS. **e**, Uni-directional tensile tests of hydrogels with and without PEDOT:PSS. **f**, 180° peeling test of the conducting hydrogel on various surfaces including polyethylene terephthalate (PET), screen printed and dried Ag ink, nitrile butadiene rubber (NBR), and mouse skin tissue. **g**, Schematic diagram illustrating the microscopic structural changes during the lower critical solution temperature (LCST) phase transition. **h**, Differential scanning calorimetry (DSC) scans of the hydrogel interface with different acrylamide (AAm) to *N-* isopropylacrylamide (NIPAM) weight ratios. The hydrogel is consisting of 150 mg mL^-1^ NIPAM, 20 mg mL^-1^ PEDOT:PSS, and AAm of 0, 9, 12, and 15 mg mL^-1^. **i,** Rheological measurement showing the phase transition temperature of the sample when the AAm to NIPAM weight ratio is 0.8:10. **j-l**, Photographs (**j**) and 180 °C peeling test (**k**) showing the drastic differences (**l**) in adhesion for the gel at room temperature and 40 °C. **m**, The tunable hydrogel adhesion can be cycled for multiple times due to the reversible nature of the LCST phenomenon. The hydrogel in **j-m** is consisting of 150 mg mL^-1^ NIPAM, 12 mg mL^-1^ AAm, and 20 mg mL^-1^ PEDOT:PSS.

Here, we designed an inter-penetrated double-network structure through *in situ* radical polymerization of a thermal-responsive covalent network of *N*-isopropylacrylamide (NIPAM)^28^ and acrylamide (AAm) in the presence of a physically crosslinked conducting polymer network of PEDOT:PSS (**Fig. 3b**). Notably, since PEDOT:PSS exists in the form of a colloidal aqueous suspension, it would severely coagulate when mixed with conventional radical initiators that contain both ionic and basic species, i.e., ammonium persulfate (AP) and *N,N,N’,N’*-tetramethylethylenediamine (TEMED) (**Fig. S6**). To ensure uniform gel formation, we developed a new initiation system based on a non-ionic redox pair of hydrogen peroxide and ascorbic acid, that allows for rapid and homogeneous gelation at room temperature (~3 min) (**Fig. S6**).

Compared to the pristine poly(NIPAM-*ran*-AAm) gel, the incorporation of PEDOT:PSS substantially reduced the interfacial impedance when in contact with phosphate buffered saline (PBS) with a ~0° phase angle across the entire frequency range (**Fig. 3c, Fig. S7**), corresponding to a resistive nature for the contact due to the high capacitance at low frequency range for PEDOT:PSS^23^. Similarly, when a voltage pulse was applied, the PEDOT:PSS gel showed substantially enhanced charge injection capacity when compared to the control sample (**Fig. 3d, Fig. S8**), which ensures efficient delivery of stimulus from the electronically conducting circuits to ionically conducting tissues. The low impedance and high charge injection of the hydrogel electrode can be well maintained even after 10,000 cycles of repetitive charge injections (**Fig. S9**).

In addition to improved electrical performances, the incorporation of PEDOT:PSS also enhanced the mechanical properties of the hydrogel. Under a uni-directional tensile test, the composite gel can be stretched to a similar strain as the control poly-NIPAM gel (~400%) but with a higher Young’s modulus, giving rise to a higher toughness (**Fig. 3e, Fig. S10**). The composite hydrogel is elastic with reversible impedance changes upon stretching to at least 100% strain (**Fig. S11**). Finally, because of the high content of polar moieties in the NIPAM-AAm backbone, the composite hydrogel can have polar interactions in addition to van der Waals interactions with diverse surfaces, such as plastic, metal, rubber, or skin, to give its strong interfacial adhesion (**Fig. 3f, Fig. S12**).

Although hydrogels containing NIPAM and PEDOT:PSS have been previously studied^29^, a conducting hydrogel with tunable adhesion has not been reported. Poly-NIPAM is a well-known polymer that exhibits a lower critical solution temperature (LCST) in water due to the heat induced aggregation of the amphiphilic NIPAM units^28^. In our case, we observed that the LCST transition was associated with drastic changes in gel adhesion, likely because the aggregated backbones can no longer form effective bonding sites with external surfaces (**Fig. 3g**). We found that additional hydrophilic monomers of AAm can be used to tune the LCST point to higher levels (i.e., above body temperature)^30^, as indicated from differential scanning calorimetry (DSC) (**Fig. 3h**). When the mass ratio between AAm and NIPAM monomers was 0.8:10, the phase change temperature reached ~40 °C, as confirmed by both DSC and rheological measurements (**Fig. 3h-i**). When tested on metal and mouse skin, the hydrogel electrodes showed strong adhesion at room temperature or normal skin temperature, comparable to 3M™ Kind Removal Silicone Tape used to secure gauze to the skin, but completely lost its adhesion with two orders of magnitude lower interfacial energy when heated above 40 °C (**Fig. 3j-l, Figs. S13-14**). Of note, the phase transition will not occur gradually before the critical temperature, as evidenced by DSC and rheology (**Fig. 3h-i**), which prevents undesired detachment during normal operation. Finally, because the LCST process is reversible^31^, the tunable adhesion of the same hydrogel can be repeated multiple times without significant degradation of the low-temperature adhesion (**Fig. 3m**).

### Validation in pre-clinical wound models

To validate our wound care management system, we performed a series of pre-clinical evaluations to test the robustness and efficacy of our developed device. Notably, mice wearing our wireless devices were able to move freely with a similar distance traveled as mice with no device attached, demonstrating an ideal therapeutic modality for patient use: namely lightweight and untethered with cables (**Fig. 4a-b**). More importantly, our temperature and impedance sensors were able to monitor the state of the wound continuously as the mice moved freely in the cage (**Fig. 4c**). In addition, our hydrogel was biocompatible and did not initiate any sensitization or irritation after continuous contact with the skin for 15 days, demonstrating no adverse reactivity signs compared to normal skin (**Fig. S15, Table S2**).

**Figure 4.**
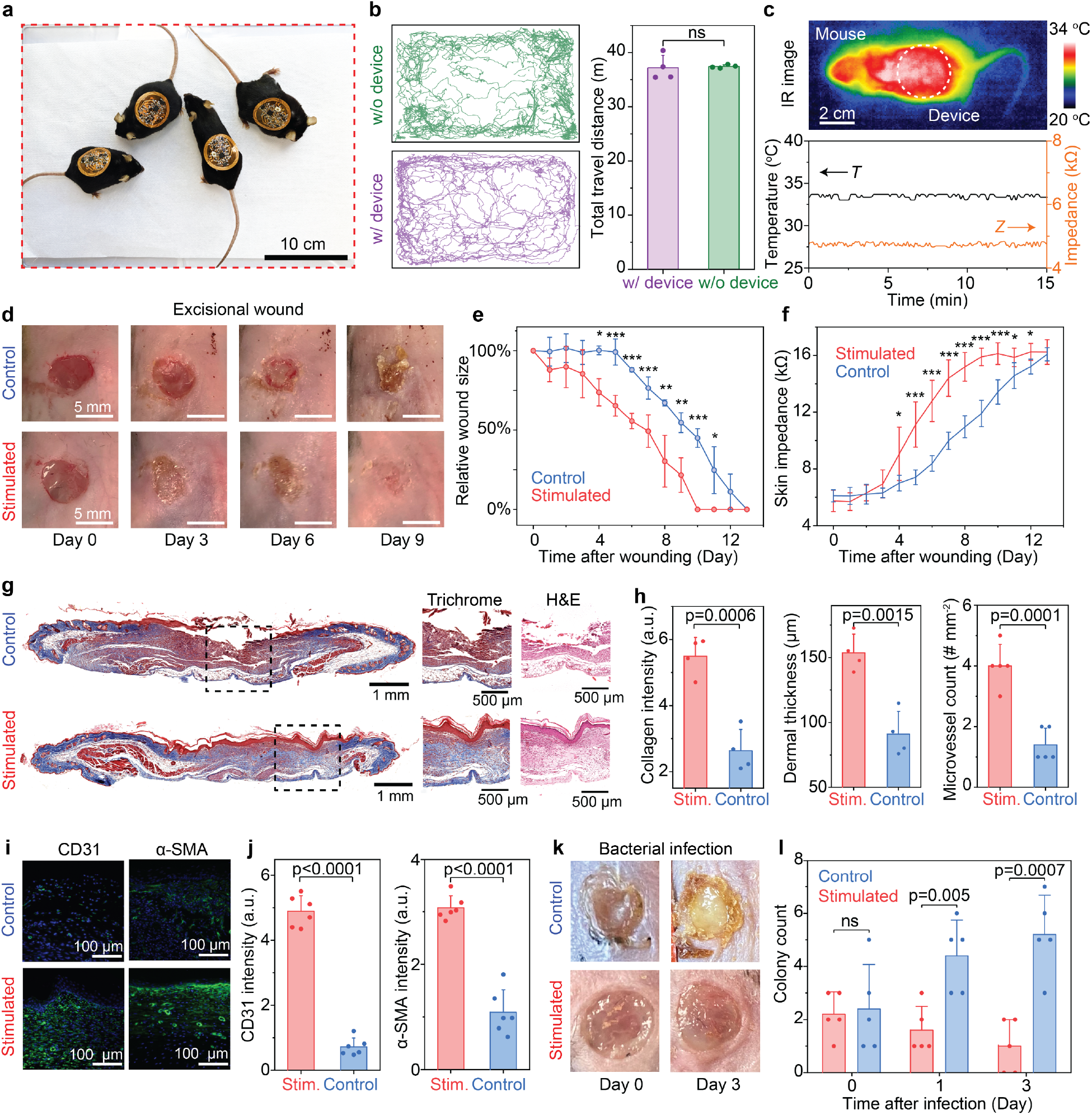
Wireless smart bandage system can continuously monitor physiological wound conditions and accelerate tissue regeneration. **a,** Photograph of four freely-moving mice wearing wireless smart bandages. **b**, Representative trajectories of mice with and without the smart bandage in the open-field test (left), and statistical analysis showing no significant differences between two groups (right). 4 mice in each group were used in the test. **c**, Infrared (IR) image of a mouse wearing the smart bandage (upper) and raw traces of wirelessly sensed wound temperature and impedance (lower). **d**, Representative photos showing the progression of wound regeneration in an excisional wound healing model with and without electrical stimulation treatment. **e** and **f**, Relative size (**e**) and impedance (**f**) of excisional wounds over time, indicating accelerated tissue regeneration with stimulation. *n*=5 for each group. All data are represented as mean ± standard deviation. Two-tailed t-test assuming equal variances were performed for the *p* values. * denotes *p* < 0.05, ** denotes *p* < 0.01, *** denotes *p* < 0.001. **g**, Representative cross-sectional histology images of skin tissue harvested from mice with and without stimulation after 13 days. Left and middle, Masson’s trichrome; Right, hematoxylin and eosin (H&E). Black dashed boxes mark the area for zoomed-in views in the middle and right panels highlighting the healed tissue. Visible intact epidermal and dermal layer observed in stimulated treatment group in the Masson’s Trichrome stain, visualized by a red surface layer which stains for muscle cells and blue layer below, which stains for collagen **h**, Quantitative comparison of collagen intensity, dermal thickness, and microvessel count for skin tissue with and without stimulation. All data are represented as mean ± standard deviation. Two-tailed t-test assuming equal variances were performed. **i** and **j**, Immunostaining images (**i**) and quantitative comparison (**j**) of CD31 and α-SMA from tissue with and without stimulation. **k** and **l**, Representative photos (**k**) and quantitative comparisons (**l**) of wounds infected with *E. coli*, with and without stimulation. All data are represented as mean ± standard deviation. Two-tailed t-test assuming equal variances were performed.

To test the device’s functionality in a biological system, a splinted excisional wound mouse model was used, where stimulated mice were treated with continuous electrical pulses. Control mice received standard sterile wound dressings without electrical stimulation. We found that stimulation resulted in accelerated wound closure (**Fig. 4d-e**) and a significant increase in wound impedance to attain a faster impedance plateau, signifying a return to an unwounded state^10,17^ (**Fig. 4f**). Stimulation of wounds also improved functional tensile recovery with increased dermal thickness, collagen deposition and overall dermal appendage count (**Fig. 4g-h, Figs. S16-18**). Of note, compared to a wired modality, our smart bandage allowed for longer and potentially continuous treatment durations (**Fig. S17**), which have been linked to accelerated wound closures^32^. Stimulated wounds also showed an increase in the collagen fiber heterogeneity, resulting in more random, shorter, and less aligned fiber orientations (**Figs. S19-20**).

We further observed a significant increase in neovascularization among stimulated wounds, with an increased microvessel count and higher CD31 and α-SMA expression (**Fig. 4h-j, Fig. S21**). Similar results were also observed in a murine burn wound healing model (**Figs. S22-25**). Our smart bandage was also found to significantly reduce infection in the wound, decreasing overall bacterial colony count (**Fig 4k-l**). Reducing infections would further improve wound care, enabling physicians to proactively treat chronic wounds, reduce hospital readmissions and medical cost, and improve patient wound healing outcomes^33^. We further validated our system in a streptozotocin (STZ) induced diabetic excisional wound model^34^, also observing an accelerated time to wound closure, improved dermal collagen fiber heterogeneity, and increased vascularization (**Figs. S26-29**). An STZ model most closely resembles Type I diabetes in patients^35^. On the cellular level, we observed the expected ability of our device to prompt cell alignment and migration, inducible with a directional electric field (**Figs. S30-31**).

### Cellular and Molecular Mechanism

Although the beneficial effects of electrical stimulation have been previously reported^20^, the cellular and molecular mechanisms for this effect remain obscure. Previous works have evaluated the role of electrical stimulation in enhancing wound healing through the activation of fibroblasts and keratinocytes, both known major cell types of the dermis that are active in the inflammatory phase of cutaneous wound repair^36–40^. Inflammatory signals activate the maturation and cross talk between these two cell types, coordinating the migration and restoration of normal tissue homeostasis after wounding^41,42^. However, the effect of electrical stimulation on immune cells, namely circulating cells, which are critical regulators of all stages of wound healing from early inflammation until late fibrosis^43,44^, remains unexplored.

We therefore decided to investigate the mechanism behind the beneficial effects of electrical stimulation on wound healing and chose to focus on the cells that infiltrate the wound from the circulation, by evaluating their transcriptional profiles using single-cell RNA sequencing (scRNA-seq). To do this, we performed parabiosis^45^ of five green fluorescence protein (GFP) positive mice to wild type (WT) mice. WT mice were wounded and either subjected to electrical stimulation or left untreated. Wound tissue from both groups was explanted on Day 5 and analyzed by scRNA-seq using the 10x Genomics Chromium platform (**Fig. 5a**). Of note, our wireless smart bandage allowed for continuous long-term stimulation, enabling the investigation of circulating cells involved in wound repair using the parabiosis model, whereas a conventional wired modality under anesthesia would not be feasible with parabiosis.

**Figure 5.**
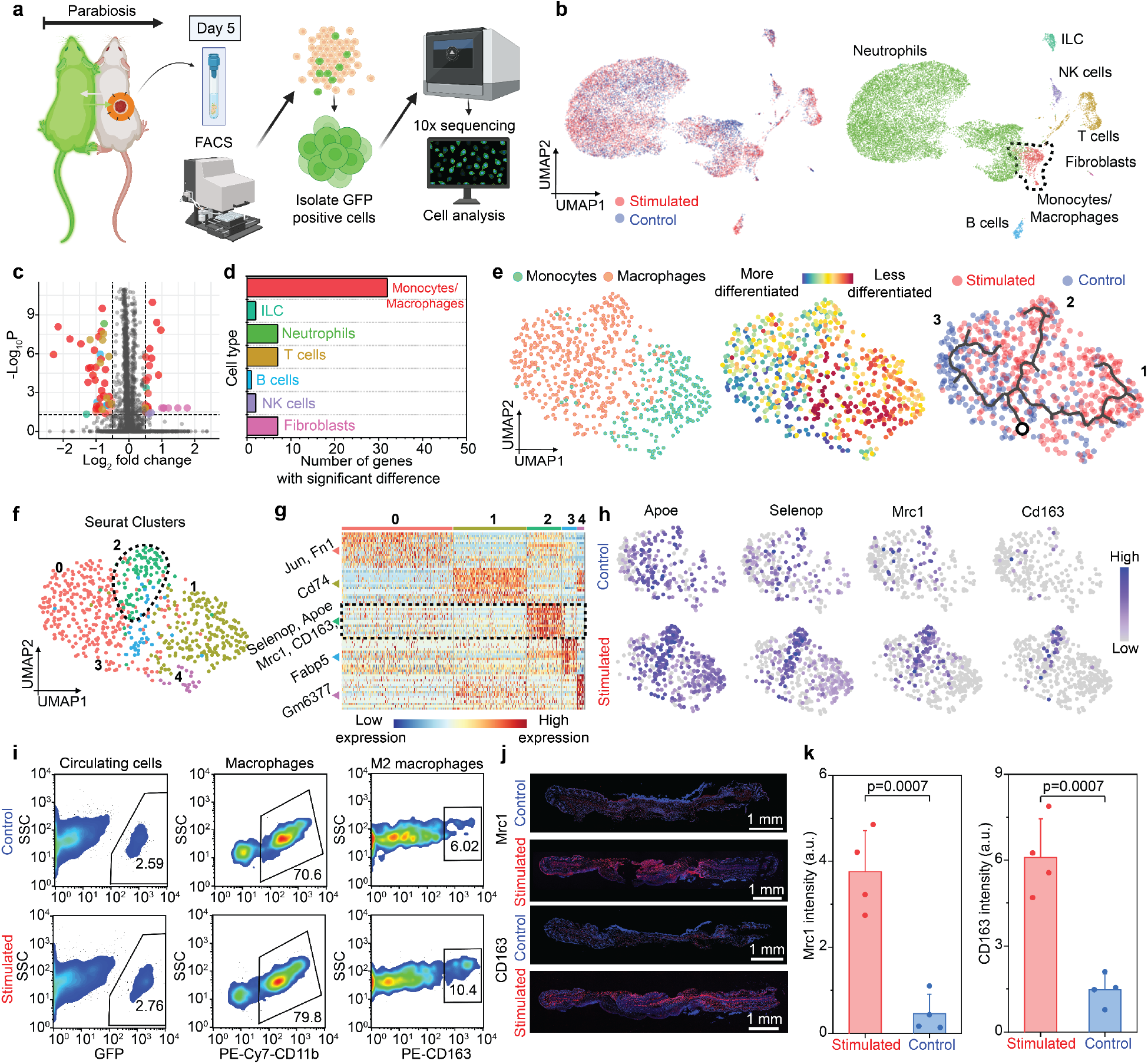
Molecular mechanism attributing to the accelerated tissue regeneration with electrical stimulation. **a,** Schematic diagram illustrating the experimental flow for the single-cell RNA sequencing (scRNA-seq). Tissues from an excisional wound of a wild type (WT) mouse paired with a GFP-positive mouse, subjected to either treatment (i.e., stimulation) or not (i.e., control), were sorted for GFP-positive cells using fluorescent activated cell sorting (FACS) and analyzed using 10x sequencing. **b,** Uniform manifold approximation and projection (UMAP) embedding of all cells colored by cell type suggesting equal overlap of stimulated and control cells. **c** and **d,** Number of differentially expressed genes (log2 fold change > 0.5 and *p* value < 0.05) for all cell types shows that the monocyte and macrophage subset have the highest number of differentially expressed genes in stimulated and untreated wounds. **e,** UMAP embedding split by macrophages and monocytes (left) verified with the CytoTrace platform (middle), which identifies differentiated cell states within the monocyte cluster. RNA velocity (right), shown as the main gene-averaged flow, visualized by velocity streamlines projected onto the UMAP embedding of the monocyte cluster categorized by treatment group and labeled with three trajectories identified by the program (trajectory 1 (right) and 2 (middle), whereas trajectory 3 (left)). These data show that monocytes are less differentiated than macrophages, as expected. RNA velocity using Monocle 3 suggests three potential fates for macrophages and monocytes, starting with the initial node, marked with a circle in the right panel. Three distinct trajectories were observed, with stimulated cells clustering along trajectory 1 and 2, while trajectory 3 was mainly composed of unstimulated cells. **f,** UMAP embedding of macrophage and monocyte Seurat clusters, grouped by cells of similar differential expression, with a pro-regenerative cluster 2, outlined with a dotted black circle, shows that there are five transcriptionally distinct clusters. Cluster 0 consists mainly of macrophages and unstimulated cells. Clusters 1, 2 and 3 consist of stimulated monocytes and macrophages. **g,** Heatmap of the top differentially expressed genes in each Seurat cluster in **f** show cluster 0 has a higher expression of genes associated with wound healing, whereas cluster 1, 2, and 3 have a higher expression of genes involved in the wound repair process. **h,** Feature plots, split by treatment (stimulated) and control, of differentially expressed genes upregulated in cluster 2 in the macrophages and monocytes indicate that there is an enrichment for pro-regenerative markers localized around cluster 2 and trajectory 2, consisting primarily of stimulated macrophages. **i,** FACS plots for treatment and control groups of GFP-positive cells circulating in the parabiosis wound model verify a higher percentage of M2 macrophages in the stimulated group. **j, k,** and quantitative comparison (**j**) of Mrc1 and CD163 from tissue with and without stimulation verifying M2 macrophage markers.

Of all the circulating inflammatory cells that were identified (**Fig. 5b, Fig. S32**), monocytes and macrophages had the highest number of differentially expressed genes in electrically stimulated and untreated wounds (**Fig. 5c, d, Fig S33**). Even with many neutrophils present, the magnitude of differentially expressed genes did not reach statistical significance (**Fig. S32, S33**). Similarly, while there were a higher number of B and T cells in the stimulated group, signifying greater recruitment of these cells from the circulation^46,47^, the overall number of cells was low and the amount of differentially expressed genes was nominal (**Fig. S32, Fig S33**).

To specifically investigate the macrophages and monocytes, we performed a series of evaluations to validate and define the high number of differentially expressed genes observed. First, we re-embedded our macrophages and monocytes and used CytoTRACE to confirm that our defined monocytes possessed less differentiated cell states based on the distribution of unique mRNA transcripts (**Fig. 5e**). We then overlaid the stimulated and unstimulated macrophages and monocytes and performed RNA velocity and pseudotime analyses using scVelo and Monocle 3, respectively, to combine RNA velocity information with trajectory inference to compute a map of potential fates that the macrophages and monocytes can undertake in response to electrical stimulation. We first used scVelo to infer our root node and transcriptional directionality across the manifold based on mRNA splicing of the macrophages and monocytes. We found three general transcriptional vector paths in which mRNA splicing could occur within individual cells, with a relatively higher amount of differentiated individual cells found on the left of the embedding and less differentiated cells found on the right (**Fig. S34a**), further confirming CytoTRACE. We then performed pseudotime analysis using Monocle 3, using a root node identified with scVelo (marked with a circle in the right panel of **Fig. 5e**) to infer terminal cell states^48^. Our analysis once again revealed 3 distinct transcriptional trajectories, with stimulated cells clustered mainly along trajectory 1 (right) and 2 (middle), while trajectory 3 (left) was mainly composed of unstimulated cells (**Fig. 5e, Fig. S34b**).

To further understand why the macrophages and monocytes had a higher amount of differentially expressed genes, we performed uniform manifold approximation and projection (UMAP) based clustering which revealed 5 transcriptionally distinct clusters (**Fig. 5f**). Of the five clusters, cluster 0, consisting of both macrophages and unstimulated control cells, had a higher expression of genes such as *Jun* and *Fn1*^49,50^, which have previously been associated with wound healing, whereas clusters 1, 2, and 3, consisting predominantly of stimulated monocytes and macrophages, demonstrated elevated expression of genes that are involved in the wound repair process, such as *Cd74, Selenop, Apoe, Mrc1, Cd163*, and *Fabp5*^51–54^ (**Fig. 5g**).

Interestingly, when we looked at the stimulated and control feature plots of highly expressed genes in macrophages and monocytes, we saw that cells with a strong enrichment for pro-regenerative markers, notably *Cd163* and *Mrc1* (CD206), as well as *Selenop* and *Apoe*, all localized around Seurat cluster 2 and trajectory 2 (middle), which primarily contained stimulated macrophages (**Fig. 5h, Fig. S36).** *Cd163* and *Mrc1* (CD206) have been previously described as M2 anti-inflammatory macrophage markers^55^, and *Selenop* has been found to be anti-inflammatory, regulating macrophage invasiveness and other inflammatory mediators responsible for pathogen clearance and tissue repair, and is linked to M2-marcophage markers such as *Stab1, Sepp1* and *Arg1*^52^. *Apoe* has been also shown to enhance *in vitro* phagocytosis of macrophages, increasing muscle and soft tissue regeneration^56,57^.

We further confirmed these transcriptional changes on the protein level, performing flow cytometry on GFP-positive cells circulating to wounds in our parabiosis model. We identified a higher percentage of CD163-positive cells in stimulated wounds as compared to controls (**Fig. 5i, Fig. S37**). This was further confirmed by immunofluorescent staining of healed tissue, with significantly higher CD163 and CD206 expression observed in stimulated wounds as compared to untreated wounds (**Fig. 5j-k**).

These data suggest that electrical stimulation may drive macrophages towards a more regenerative phenotype and could underly the accelerated wound healing observed in our pre-clinical studies. The high predominance of regenerative macrophages could be in part due to macrophages responding to the local microenvironmental stimuli. Modulating the cell membrane electric potential with electrical stimuli could activate more KATP ion channels, which has previously been shown to affect macrophage differentiation plasticity and function^58,59^. Taken together, our pre-clinical studies identify one mechanism by which electrical stimulation contributes to the coordination and regulation of macrophage functions, including those essential for microbial clearance and wound healing.

### Conclusions

In summary, we designed and fabricated a miniaturized smart bandage with dual channel continuous sensing of wound impedance and temperature, as well as a parallel stimulation circuit to deliver programmed electrical cues for accelerated wound healing. Our wireless smart bandage system can provide: (i) active monitoring and continuous treatment of the wound, and (ii) accelerated healing through a pro-regenerative mode of action, activated by continuous electrical stimulation across the wound bed that increases cellular proliferation, activation, and recruitment of cells involved in wound repair. With further integration of other on-board sensors, actuators and computational modules, our wireless system can also be adapted to other disease management, allowing for the next generation of closed-loop bioelectronic medicine.

## Acknowledgments

This work was supported by the Stanford Clinical and Translational Science Award (CTSA) to Spectrum. The CTSA program is led by the National Center for Advancing Translational Sciences (NCATS) at the National Institutes of Health (NIH). This work was partly supported by BOE Technology Group Co., Ltd. Part of this work was performed at the Stanford Nano Shared Facilities (SNSF), supported by the National Science Foundation under award ECCS-2026822. We thank Agfa for providing PEDOT:PSS. We thank Theresa Carlomagno and Tiffine Vang for administrative support. We thank Yu Jin Park for tissue histology support, Doreen Wu in the Stanford Animal Histology Services and Pauline Chu in the Human Research Histology Core for help with preparation of histologic specimens. We thank Siavash Kananian for instrument support of VNA measurements. We also thank Dr. Russ Altman for his guidance with the project.

## Author Contributions

Y.J., A.A.T., S.N., G.C.G. and Z.B. designed the study. S.N., Y.J. performed circuit design and testing. Y.J., C.-C.S., J.-C.L., D.Z. J.T. performed material synthesis and characterizations. A.A.T., Y.J., D.H., K.C., A.M.M, S.M., M.R.L., A.S., E.B., S.J., S.R.S., K.S., T.J., E.Z., C.R.N., W.G.V., D.S., J.P., M.R., D.P.P., A.C., M.C.L., C.A.B., S.H.K., K.S.F., G.G. K.L., K.Z. performed the animal, cell culture experiments and single cell evaluations. Y.J., A.A.T., S.N., M.J., G.C.G. and Z.B. wrote the manuscript with input from all co-authors.

## Methods

A complete, detailed description of methods can be found in the Supplementary Information.

