## Supplementary Information for "Wireless closed-loop smart bandage for chronic wound management and accelerated tissue regeneration"

### Table of Contents

**Table S1 | Comparison of currently available smart bandage technologies and key differentiators to our work. 12**
**Figure S1 | Loading curve of the antenna wireless energy harvesting. 13**
**Figure S2 | Validation of the high-pass filter (HPF) design for impedance sensing. 14**
**Figure S3 | Antenna characterization under bending. 15**
**Figure S4 | Validation of the sensing performance under simultaneous stimulation. 16**
**Figure S5 | Closed-loop operation of the wireless smart bandage. 17**
**Figure S6 | Comparison of the gelation outcome using different initiators. 18**
**Figure S7 | EIS measurement (a) and electrical conductivity (b) of the hydrogel electrode with different concentrations of PEDOT:PSS. 19**
**Figure S8 | Chronoamperometry measurement (a) and injected current (b) of the hydrogel electrode with different concentrations of PEDOT:PSS. 20**
**Figure S9 | Cyclic tests of EIS (a) and chronoamperometry (b) of the hydrogel electrode showing the high electrochemical stability after at least 10000 cycles of repetitive charge injections. 21**
**Figure S10 | Uni-directional tensile test of the composite hydrogel electrode. 22**
**Figure S11 | Cyclic loading of the hydrogel electrode and corresponding impedance changes. 23**
**Figure S12 | 180-degree peeling test of the hydrogel adhesion onto various substrates. 24**
**Figure S13 | Photos showing the tunable adhesion of the same hydrogel electrode with a hot plate under different temperature. 25**
**Figure S14 | Photos showing the tunable adhesion of the same hydrogel electrode with moust skin under different temperature. 26**
**Table S2 | Semi-quantitative scoring criteria for paraffin sections to assess the hydrogel biocompatibility. 27**
**Figure S15 | Hydrogel electrode showing excellent biocompatibility after long-term contact with mouse skin. 28**
**Figure S16 | Electrical stimulation induces higher skin toughness after healing compared to the control tissue. 29**
**Figure S17 | Wireless smart bandage allows longer treatment per day with better healing outcomes. 30**
**Figure S18 | Histological studies show improved wound healing outcomes for excisional wounds treated with electrical stimulation. 31**
**Figure S19 | FracLac evaluation shows less scar-like fiber arrangement for excisional wounds treated with electrical stimulation after 13 days. 32**
**Figure S20 | Picrosirius red staining shows more random collagen networks for excisional wounds treated with electrical stimulation after 13 days. 33**
**Figure S21 | Histology and immunostaining reveals more neovascularization for excisional wounds treated with electrical stimulation after 13 days. 34**
**Figure S22 | Histological studies show improved healing outcomes for burn wounds treated with electrical stimulation after 19 days. 35**
**Figure S23 | FracLac evaluation shows less scar-like fiber arrangement for burn wounds treated with electrical stimulation after 19 days. 36**

#### Methods

##### Fabrication of the flexible circuit

Flexible printed circuit boards (FPCB) with custom designs are manufactured by commercial vendors with ISO 9001 certificate, e.g, PCBWay. The bill of materials includes passive components (capacitors, resistors, and diodes), a thermistor (103KT1608T-1P, Semitec), a near-field communication (NFC) ISO15693 sensor transponder with a programmable low-power microcontroller (MCU, RF430FRL152H, Texas Instruments), a crystal oscillator (SG-3040LC 32.7680KB3: PURE SN, Epson Timing), and two operational amplifiers (TSV620AILT, STMicroelectronics). All circuit components were soldered using tin-lead solder paste (Sn63/Pb37, melting point: 183 ºC, 247 Solder) by hot-air blowing.

To avoid chronic electrochemical corrosion of the electroless nickel immersion gold (ENIG) electrodes on the backside of the smart bandage board (two for stimulation, two for sensing), silver/silver chloride (Ag/AgCl) conducting paste (CI-4040, Nagase ChemteX) was printed onto each electrodes using blades and cured on hot plate at 100 ºC for 10 min. The electrode area was further encapsulated by an elastomeric polyurethane coating (Clear Flex 50, Mix Ratio: 1A/2B wt/wt, Smooth-On) and cured at 70 ºC for 15 min. The hydrogel interface electrodes were then placed onto corresponding Ag/AgCl electrodes for sensing and stimulation purposes.

The resonant frequency and quality factor of the antenna was measured using a portable vector network analyzer (VNA, miniVNA Tiny Plus 2). The resonant frequency was determined by the peak location of the impedance amplitude. The quality factor was calculated by resonant frequency divided by the 3 dB bandwidth. Wireless powering and readout were performed using a 13.56 MHz desktop reader/writer with built-in antenna (Gao RFID). A custom written software was used to control the powering pattern and continuous readout.

##### Synthesis of conducting adhesive hydrogel

In a typical synthesis, 40 mg of poly(3,4-ethylenedioxythiophene) polystyrene sulfonate dry pellet (Orgacon™ DRY5, Agfa), 300 mg of *N*-isopropylacrylamide (NIPAM, Sigma-Aldrich), 24 mg of acrylamide (AAm, Sigma-Aldrich), and 1 mg of *N,N'*-methylenebisacrylamide (MBAA, Sigma-Aldrich) were mixed and dissolved in 2 mL of water by stirring for 30 min at room temperature. Prior to the gel initiation, 100 µL of 30% (v/v) hydrogen peroxide ($H_2O_2$, Fisher Scientific) and 100 µL of 20% (wt/v) ascorbic acid (AA, Sigma-Aldrich) were added sequentially to the monomer mixture and quickly mixed before pouring into Teflon molds for subsequent gelation at room temperature.

For the control sample without PEDOT:PSS, the same procedure was performed except for the absence of PEDOT:PSS pellet in the initial mixture. For the study of AAm content on the overall lower critical solution temperature (LCST) behavior, the initial amount of AAm was varied from 0 mg, 18 mg, 24 mg, and 30 mg. For the study of electrochemical properties, the initial amount of PEDOT:PSS was varied from 0 mg, 10 mg, 20 mg, 30 mg, and 40 mg. For the study of crosslinking density, the initial amount of MBAA was varied from 0.3 mg, 1 mg, 3 mg, and 10 mg.

##### Electrochemical characterizations

Electrochemical impedance spectroscopy (EIS) was performed by using the hydrogel as the working electrode, platinum as the counter electrode, silver/silver chloride (Ag/AgCl) as the reference electrode, and PBS as the electrolyte. Impedance and phase angle as functions of frequency were acquired by a Bio-Logic VSP-300 workstation with a sine wave signal amplitude of 10 mV. Chronoamperometry was performed on the same potentiostat by delivering biphasic square wave pulses with 50 ms for each phase and 100 mV amplitudes with currents being recorded simultaneously.

Conductivity measurements were carried out using a four-point probe method using a Keithley 4200 SC semiconductor analyzer. For the measurement of impedance change over strain, the conducting hydrogel was mounted onto a home-made automated stretcher when the impedance values at different strain levels would be recorded using a LCR meter (Keysight Technologies, E4098A).

#### Mechanical characterizations

Uni-directional tensile tests of the hydrogel electrodes were performed on an Instron 5565 at a strain rate of 10 mm $min^{-1}$. To measure the interfacial energy, hydrogel samples and surfaces for adhesion (e.g., metal, plastic, rubber, and skin) were glued onto Kapton polyimide films using cyanoacrylate (Krazy Glue) as a stiff backing. After adhering the hydrogel onto the surface of interest, the adhesion was tested by the standard 180-degree peel test with the Instron machine. All tests were conducted with a constant peeling speed of 10 mm $min^{-1}$. Interfacial toughness was calculated by dividing two times the plateau by the width of the tissue sample. Differential scanning calorimetry (DSC) was conducted using a TA Instruments Q2000 DSC. Rheological measurements were performed using a TA Instruments ARES-G2 rheometer.

#### Hydrogel biocompatibility assay

All tissue sample histology slides were examined via light microscopy by the study pathologist. Wound sites were semi-quantitatively scored per the criteria of **Table S2**.

Semi-quantitative scores were assigned based on the representative site response observed over six noncontiguous representative high-powered microscope fields at the host/test sample interface. The Test (with gel) Sample and Control Sample scored results are documented by animal, as seen in **Fig. S15.** Sample Reactivity Score calculations are documented and categorized. Representative low and high magnification implant site photomicrographs are contained in **Fig. S15.** After scoring of each implant site sample or subsample slide section, the following steps were performed. Polymorphonuclear cells, lymphocytes, plasma cells, macrophages, giant cells, and necrosis scores assigned for each implant were totaled and multiplied by 2 (Subtotal A). Neovascularization, fibrosis, and fatty infiltration scores assigned for each implant were totaled (Subtotal B). Subtotal A and Subtotal B were added together for a total implant score (Total). For each Test or Control Sample type, the Total values were added together (Group Total). Each Group Total was averaged (Group Average) by the appropriate number of scored implants. The Control Group Average was subtracted from the Test Group Average to derive the final average test result (Test Sample Relative Score). A resulting negative difference was recorded as zero, per protocol. The Test Sample Relative Score, per part c in **Fig. S15,** was used to categorize the Test Sample.

#### Animal testing

##### Animals

8–12-week-old mice (eC57BL/6J; Jackson Laboratory, Bar Harbor, ME, http://www.jax.org) were housed in the Stanford University Veterinary Service Center in accordance with NIH and institution-approved animal care guidelines. Mice within one experiment were of the same strain, batch and birthday. All procedures were approved by the Stanford Administrative Panel on Laboratory Animal Care.

Open-field movement test

The open-field test for mice with and without the smart bandage was performed in a 25×40 $cm^2$ open cage. Four mice for each group were allowed to freely explore the open-field enclosure for 15 min under ambient conditions, while being recorded by an overheard camera. The recorded videos were analyzed using the DeepLabCut deep learning architecture to measure the total travel distances for each group.

Excisional splinted wound model

Twelve-week-old C57BL/6 mice (Jackson Laboratories) were randomized into the following groups: (1) stimulated, (2) non-stimulated control (*n*=5). Excisional wounds on the dorsum of the mice were created as previously described[1] using 6 mm biopsy punches. Wounds were treated daily until closure. All wounds were covered with an occlusive dressing (Tegaderm, 3M, MN). Digital photographs of the wounds were taken every other day until closure. Wound closure was defined as the time at which the wound was completely re-epithelialized without any scab. Wound area was determined using ImageJ software by a blinded observer (NIH, Bethesda, MD).

Contact burn model

An established partial thickness contact burn model was used[2]. Briefly, mice were anesthetized using 2% isoflurane and administered 0.05 mg $kg^{-1}$ buprenorphine subcutaneously. Aluminum cylinders with a 10 mm diameter (Alfa Aesar, Ward Hill, MA) were heated in a 100°C water bath for 5 minutes and applied to the dorsa of the animals for 15 seconds, with only the weight of the rod (47.75 g) applying pressure to the skin. Two burns were created on each mouse. Mice were immediately placed supine in a cool water bath for 1 minute to quench the burn. Wounds were mechanically debrided on post-burn day 5 using a blunt stainless-steel rod and randomized to treatment with stimulation, or no treatment (no stimulation) (*n*=4 for each group). Treatment was administered daily. Burns were photographed at regular intervals until completely healed and analyzed using ImageJ software (NIH, Bethesda, MD). Wound area (as percent of original) was plotted against post burn day with area under the curve.

Streptozotocin (STZ) induced wound model

To model Type I diabetic wound healing, an STZ-induced mouse model was used[3]. Eight-week-old C57BL/6 mice were fed with high-fat and high-sugar feed and maintained under specific pathogen–free environment. Briefly, STZ (100 mg $kg^{-1}$; Sigma-Aldrich) was mixed in sodium citrate buffer (intraperitoneal injection) and administered daily for two consecutive days (*n*=4 for each group, stimulated and non-stimulated control). All of the mice with plasma glucose levels ≥16.7 mM under normal condition were considered diabetic 2 weeks after the first STZ injection[4]. Animals were maintained in a diabetic state for the duration of the experiment.

Infection model

To validate the effectiveness of electrical stimulation at reducing biofilm and wound infection, an *Escherichia coli (E. coli)* wound infection model was used[5]. Briefly, 30 µL of fresh culture of *E. coli* DH5α were inoculated overnight in 10 mL lysogeny broth (LB) medium. *E. coli* was grown at 37 ºC with shaking to an O.D. of 600. After overnight incubation, tubes were vortexed, and the volume adjusted to obtain $10^7$ CFU $mL^{-1}$. Bacteria were centrifuged and re-suspended in 20 mL sterile PBS. To control the concentration of bacteria a serial dilution of the utilized suspension was prepared and inoculated on fresh LB agar plate followed by incubation for 18 h at 37°C. Colonies were counted on the next day to determine the original bacteria concentration.

On the day of inoculation, 6 mm excisional wounds were created on the dorsum of the mouse and inoculated with 10 μL of PBS containing $10^7$ CFU bacteria using a pipette tip. Tegaderm (3M, MN) were cut and applied to the wound area. Wounds were treated with stimulation or left untreated. Immediately after treatment, the wound and wound periphery were swabbed with a BD ESwab Collection & Transport System (BD Biosciences, CA) and submitted to the Stanford Diagnostic Lab for quantification. An *n*=5 was submitted for each treatment group at time points 0 h, 24 h, and 72 h post inoculation.

A semi quantitative method was used to count total bacterial colony count. A sample was plated from the ESwab fluid, then spread into quadrants. If the bacteria grew where the original specimen was plated that is referred to as 1+ growth (or very light colony growth), 2+ extends to the edges of the first quadrant and outside the 1+ region (light colony growth), 3+ is considered moderate colony growth and is colony growth that extends through the 3rd quadrant, 4+ is heavy colony growth and grows into the 4th quadrant. Data was subsequently aggregated and analyzed. Digital photographs of the wounds were taken every day of specimen collection.

Parabiosis model

To study the role of cell migration into the wound, a parabiosis model was used as previously described[6]. Briefly, the corresponding flanks of $GFP^+$ (C57BL/6-Tg(CAG-EGFP)1Osb/J) and C57BL6/J mice (WT) were shaved and disinfected with Betadine solution and 70% ethanol three times. Matching skin incisions were made from the olecranon to the knee joint of each mouse. The skin edges were undermined to create skin flaps of 1 cm width. 6-0 nylon sutures (Ethilon) were used to approximate the dorsal and ventral edges of the skin flaps and skin staples were used to close the longitudinal incisions. Buprenorphine was used for analgesia by subcutaneous injection every 8–12 h for 48 h after operation. Mice were monitored daily until the end of the experiment. After 14 days, peripheral blood chimerism was confirmed using fluorescent microscopy of the tail vein blood. After cross-circulation is established between the two parabionts, an excisional splinted wound is created on the dorsum of the WT mouse. Wounds are subsequently treated with stimulation or left untreated. *n*=5 for each group.

On day 5, analysis of murine cells isolated from excisional wounds was performed according to published protocols for flow cytometry on murine tissue[7]. Briefly, tissue was explanted on day 5 from the parabiosis model, then micro-dissected and incubated in serum-free Dulbecco's Modified Eagle Medium (DMEM) with 240 U of collagenase IV per mL for 1h at 37°C in a rotating oven. Digested tissue was filtered, centrifuged and stained with PE/Cy7 rat anti-mouse CD11b antibody, PE rat anti-mouse CD163 antibody, and APC rat anti-mouse F4/80 antibody (Biolegend, San Diego, CA). 4′,6-diamidino-2-phenylindole (DAPI) was used to stain dead cells. Flow cytometry was performed on a BD FACS Aria (Becton Dickinson, San Jose, CA) and data was analyzed using FlowJo (Becton Dickinson, San Jose, CA). GFP+ cells from stimulated and non-stimulated control groups were sorted into collection tubes and submitted for 10x sequencing.

Skin impedance and temperature sensing

The smart bandage was attached to the mouse skin interfaced by the hydrogel electrodes. Continuous wireless readout of temperature and impedance was performed using a 13.56 MHz desktop reader/writer with built-in antenna (233015, Gao RFID). A custom written software was used to control the readout sampling interval and data storage.

An infrared (IR) thermal imaging camera (HT-19, Hti-Xintai) was used to capture thermal mouse images.

#### Tissue characterization

##### Histology and staining

Wounds were harvested with a 2-mm rim of unwounded skin from euthanized mice. Skin tissues were fixed overnight in 4% paraformaldehyde followed by serial dehydration in ethanol and embedding in paraffin as previously described[1]. 5 µm sections (*n*=5) were stained with hemolysin and eosin (H&E) and Masson's trichrome (Sigma-Aldrich). Nuclei were stained with Hoechst dye. Tissue was stained for picrosirius red (Sigma-Aldrich). For CD31 (ab28364, Abcam), α-SMA (ab5694, Abcam), CD206 (ab64693, Abcam), and CD163 (ab182422, Abcam) DAPI (ab104139, Abcam) was used to stain for nuclei.

For immunohistochemical staining, slides were first deparaffinized in xylene, and then rehydrated in ethanol/PBS mixtures. The slides were then washed three times in PBS. Next the slices underwent antigen retrieval using 1× sodium citrate pH 6.0 (100× diluted in PBS; Abcam) in deionized (DI) $H_2O$. The slices, submerged in the solution, were warmed in a microwave on full power for 90 seconds. One minute later, at 60% power for 60 seconds. One minute later, the slices in solution were places in 4 °C for 30 minutes and subsequently soaked in DI $H_2O$ for 5 minutes. The slices were then washed three times in PBS.

Slices were permeabilized for intracellular antigens with 0.2% Trion X-100 (Sigma-Aldrich) for 10 minutes, then washed three times with PBS. A PAP/hydrophobic pen was used to isolate slices. Slices were then blocked with a solution consisting of 5% goat serum (vol/vol; Sigma-Aldrich) in PBS for 2 hours at room temperature in a humidified chamber.

Slices were then incubated for 1 day at 4 °C with primary antibodies in blocking solution. Sections were then washed three times with PBS for 30 min and then stained for 2h at 4 °C with corresponding secondary antibodies (Goat anti-Mouse IgG (H+L) Highly Cross-Adsorbed Secondary Antibody, Alexa Fluor Plus 488, A32723, 1:250, Thermo-Fisher; Goat anti-Rabbit IgG (H+L) Highly Cross-Adsorbed Secondary Antibody, Alexa Fluor Plus 647, A32733, 1:250, Thermo-Fisher). Slices were washed three times with PBS and incubated with DAPI (4′6-diamidino-2-phenylindole) (1:50,000) for 30 min.

H&E and Masson's trichrome were imaged with brightfield microscopy. Picrosirius red imaged using polarized light microscopy (Leica DM5000 B upright microscope) to obtain 40× magnification images. CD31, α-SMA, CD206, and CD163 were imaged using a Zeiss LSM 880 confocal laser scanning microscope at the Cell Science and Imaging Facility at Stanford University. To obtain high resolution images of tissue samples, multiple images of 25x magnification were acquired using automatic tile scanning and were stitched together using Leica LAS-X software.

##### Tissue analysis

Image J (NIH, Bethesda, MD) was used to binarize images taken with the same settings, and intensity thresholds were used to quantify staining based upon pixel-positive area averaged over three high power fields. All histology and immunofluorescent images shown are representative images of multiple experiments and were performed by a blinded observer. Dermal thickness was quantified using Image J, where the distance from the epidermis to the dermis, visible in red with Masson's trichrome, was measured. Luminal structures, visible with Masson's trichrome staining, containing red blood cells were considered dermal microvessels. Three high power fields at 40x were examined for each wound sample to quantify microvessels. Appendages were quantified using Masson's trichrome with three high power fields at 5×.

Dermal collagen analysis

Picrosirius red images were used for the following analysis. The CT-FIRE algorithm was used to analyze individual fiber metrics such as length, width, angle, and curvature. It can also extract other variables such as localized fiber density and the spatial relationship between fiber and the associated boundary[8]. CurveAlign 4.0 was used to quantify all fiber angles and strength of alignment within an image[9]. Complexity and heterogeneity were measured using the ImageJ plug-in FracLac[10]. Local fractal dimensions (FD) and lacunarity (L) values were calculated using the subsample box counting scan (50 grid default sampling size, minimum pixel density threshold = 0, rectangle subscan). FD measures density of collagen networks. A higher FD has a denser and scar-like fiber arrangement. L measures the amount of randomness or heterogeneity in a sample. A low L implies less heterogenous collagen fiber orientation. Finally, MatFiber was used to further quantify fiber alignment, reported as mean vector length. The strength of alignment ranges from a value of 0 (completely random fiber alignment) to 1 (completely aligned fibers)[11]. The average fiber parameters for each mouse were used for statistical analysis.

Wound tensile testing

Biomechanical testing was performed as previously described[12]. Briefly, uniaxial tensile testing was performed at room temperature using a tensile testing apparatus (Bionix 200; MTS Systems Corporation, Eden Prairie, MN). Constant volume deformation was assumed during uniaxial stretching.

***In vitro* study of cell alignment and migration**

Human umbilical vein endothelial cells (HUVEC, Sigma-Aldrich) were cultured on glass-bottomed Petri dishes and passaged following standard procedures from the vendor.

For the alignment/migration study, a silicone spacer based on polydimethylsiloxane (PDMS) (25 mm×5 mm×10 mm, L×W×H) was glued onto a glass bottomed dish using a biocompatible silicone adhesive (Kwik-Sil, World Precision Instruments) with a small opening at the bottom (5 mm×5 mm×200 μm, L×W×H) to allow medium exchange and establish the electric field. HUVEC cells were seeded in the opening of the spacer to allow cell to adhere and proliferate for 24 hours. The cells were then stained with 2 μM of Calcein AM (Thermo Fisher) for 30 min at 37 °C and washed three times with dye-free culture media before imaging.

During the experiment, using an agar salt bridge, a waveform mimicking wireless stimulation condition was applied with a 13.56 MHz sinusoidal wave of 1 V peak-to-peak amplitude. Live cells were imaged in real time using a Leica TCS SP8 confocal microscope (Leica) and analyzed using ImageJ software.

**Single cell analysis**

Single cell RNA sequencing

The GFP+ cell suspension was resuspended in a concentrated solution and submitted for droplet-based microfluidic single cell RNA sequencing (scRNA-seq) at the Stanford Functional Genomics Facility (SFGF) using the 10x Chromium Single Cell platform (Single Cell 3' v3, 10x Genomics, USA). The cell suspension, reverse transcription master mix, and partitioning oil was loaded onto a single cell chip, processed on the Chromium Controller, and reverse transcription was performed at 53°C for 45min. cDNA was amplified for 12 cycles total (BioRad C1000 Touch thermocycler) with cDNA size selected using SpriSelect beads (Beckman Coulter, USA) and a 3:5 ratio of SpriSelect reagent volume to sample volume. cDNA was analyzed on an Agilent Bioanalyzer High Sensitivity DNA chip for qualitative control, fragmented for 5 min at 32 °C, followed by end repair and A-tailing at 65 °C for 30 min, and then double-sided size selected with SpriSelect. Sequencing adaptors were ligated to the cDNA at 20 °C for 15min. cDNA was amplified using a sample-specific index oligo as primer, followed by another round of double-sided size selection. Final libraries were analyzed on an Agilent Bioanalyzer High Sensitivity DNA chip for qualitative control purposes. cDNA libraries were sequenced on a HiSeq 4000 Illumina platform aiming for 50,000 reads per cell.

Single cell RNA-seq data processing, normalization, and cell cluster identification

Base calls were converted to reads using the Cell Ranger (10x Genomics, Pleasanton, CA, USA; version 3.1) implementation mkfastq and then aligned against the MM 10 (mouse) genome using Cell Ranger's count function with SC3Pv3 chemistry and 5,000 expected cells per sample. Cell barcodes representative of quality cells were delineated from barcodes of apoptotic cells or background RNA based on a threshold of having at least 300 unique transcripts profiled, less than 100,000 total transcripts, and less than 10% of their transcriptome of mitochondrial origin. Unique molecular identifiers (UMIs) from each cell barcode were retained for all downstream analysis. Raw UMI counts were normalized with a scale factor of 10,000 UMIs per cell and subsequently natural log transformed with a pseudocount of 1 using the R package Seurat (version 3.1.1)[13]. Aggregated data were then evaluated using uniform manifold approximation and projection (UMAP) analysis over the first 15 principal components[14]. Cell annotations were ascribed using the SingleR package (version 3.11) against the ImmGen database[15,16] and confirmed by expression analysis of specific cell type markers. Louvain clustering was performed using a resolution of 0.5 and 15 nearest neighbors. Cell-type markers were generated using Seurat's native FindMarkers function with a log fold change threshold of 0.25 using the ROC to assign predictive power to each gene.

Pseudotime analysis

Pseudotime analysis was performed using the Monocle 3 package in R (version 3 0.2.0)[17]. A principal graph was learned from the reduced dimension space using reversed graph embedding with default parameters. The principal GraphTest function using the Moran's I statistic was employed to identify correlated genes on trajectory embedded in the manifold.

RNA velocity analysis

RNA velocity analysis was performed using scVelo[18]. In contrast to previous steady-state models, which falsely assume that all genes share a common splicing rate, scVelo uses a likelihood-based dynamical model to solve the full transcriptional dynamics of splicing kinetics. Thereby RNA velocity analysis can be adapted to transient cell states and heterogeneous cellular subpopulations as in our dataset. Partition-based graph abstraction (PAGA) was performed using the sc.tl.paga function in scVelo. To find genes with differentially regulated transcriptional dynamics compared to all other clusters a Welch t-test with overestimated variance to be conservative was applied, using the sc.tl.rank_velocity_genes function. Genes were ranked by their likelihood obtained from the dynamical model grouped by Seurat clusters. A pseudotime heatmap was created by plotting the expression of the genes identified by sc.tl.rank_velocity_genes along velocity-inferred pseudotime. The terminal transcriptional states were identified as end points of the velocity-inferred Markov diffusion process.

<u>CytoTRACE</u>

CytoTRACE (Cellular Trajectory Reconstruction Analysis using gene Counts and Expression) was used to infer the cell differentiation trajectories[19,20]. Briefly, this algorithm places the cells along a trajectory corresponding to a biological process (cell differentiation) by taking advantage of an individual cell's asynchronous progression under an unsupervised framework.

#### **Statistical analysis**

Statistical significance was determined using a two-tailed unpaired t-test for experiments involving only two conditions. For experiments with 3 or more conditions, pairwise t-tests were employed, and *p*-values adjusted using Bonferroni correction for multiple hypothesis testing. A $p$-value $<0.05$ was considered statistically significant. The statistical methods used for scRNA-seq analysis are described in the specific sections above.

| Reference | Summary of Function | Sensing modality | Treatment modality | Wireless (Y/N) | Closed-loop (Y/N) | Biological model |
|---|---|---|---|---|---|---|
| *Adv. Healthcare Mater.* **5**, 711-719 (2016) | pH-responsive dyes incorporated into flexible hydrogel fibers for real-time pH measurement | • pH | N/A | N | N | • Ex vivo healthy tissue |
| *Adv. Mater.* **28**, 502-509 (2016) | A transparent stretchable gated sensor array with responsivity to temperature and strain changes | • Temperature<br>• Strain | N/A | N | N | • Healthy skin |
| *Adv. Healthcare Mater.* **3**, 1597-1607 (2014) | An epidermal electronics system that records temperature and thermal conductivity of the skin tissue | • Temperature<br>• Thermal conductivity | N/A | Y | N | • Healthy skin |
| *Sci. Adv.* **6**, eabd1061 (2020) | A brightly emitting phosphorescent porphyrin embedded within a paintable liquid bandage formulation for measuring tissue oxygenation | • Oxygenation | N/A | N | N | • Patients undergoing skin-sparing mastectomy |
| *Nat. Commun.* **6**, 6575 (2015) | A flexible, electronic device that non-invasively detects pressure-induced tissue damage using impedance spectroscopy across electrode arrays. | • Impedance | N/A | N | N | • Pressure ulcer |
| *Biosens. Bioelectron.* **31** 413-418 (2012) | Screen printed electrodes running electrochemical impedance spectroscopy for the detection and quantification of three infection biomarkers | • Infection biomarkers | N/A | N | N | • In vitro buffer |
| *Small* **14**, 1703509 (2018) | A drug releasing system comprising of a pH and temperature sensor as well as a hydrogel loaded with thermo-responsive drug carriers | • Temperature<br>• pH | Antibiotic drug release | N | N | • In vitro scratch assay |
| **Our work** | A flexible bioelectronic system consisting of wirelessly powered, closed-loop sensing and stimulation circuits with tissue-interfacing tough conducting hydrogel electrodes for continuous monitoring of wound temperature and impedance as well as electrotherapy | • Temperature<br>• Impedance | Electrotherapy through galvanotaxis | Y | Y | • Splinted excisional model<br>• Burning model<br>• STZ-induced diabetes model<br>• Infection model |

**Table S1 | Comparison of currently available smart bandage technologies and key differentiators to our work.**

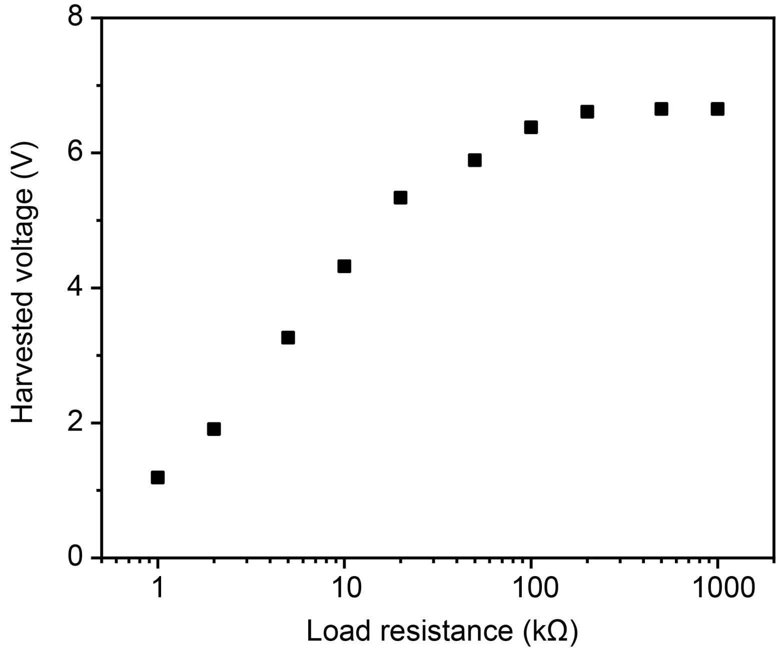

**Figure S1 | Loading curve of the antenna wireless energy harvesting.**

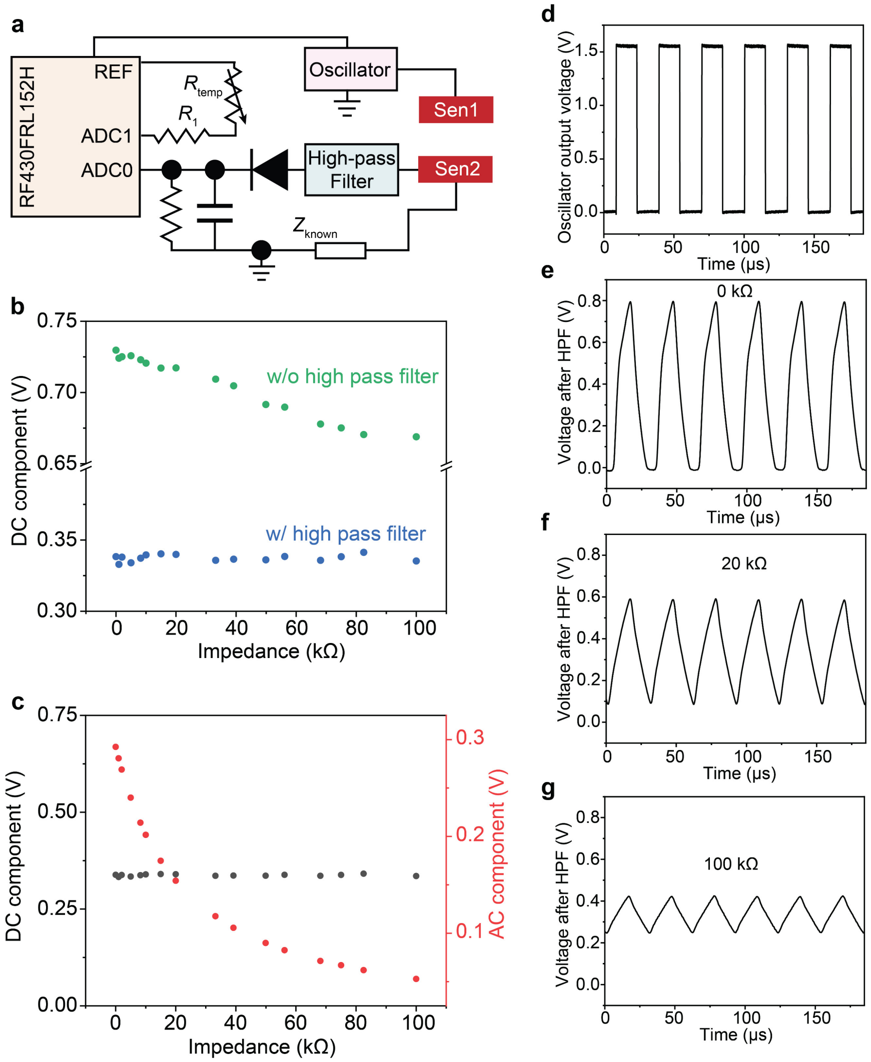

**Figure S2 | Validation of the high-pass filter (HPF) design for impedance sensing. a,** Circuit diagram showing the sensing design. For temperature sensing, the RF430FRL152H transponder has direct thermistor support (ADC1 channel) by emitting a small µA level current on the thermistor and sampling the voltage. For the impedance sensor, an oscillator was used to generate a 32.768 kHz square wave alternating current (AC) signal that passed through the wound and a known impedance component ($Z_{known}$). Through a voltage divider, the AC signal applied on $Z_{known}$ could then reflect the wound impedance. This received AC signal was further conditioned through a HPF to remove the direct current (DC) component inside the oscillation signal. Finally, an envelope detector was used to convert the AC signal amplitude to a DC voltage, which was captured by the ADC0 channel inside the RF430FRL152H transponder. **b,** Comparison of the DC component with and without the HPF. Without the HPF, the DC component would change under different load impedance, which would interfere with the AC amplitude for sensing. **c,** Voltage output after the high-pass filter showing reduced AC amplitudes with respect to larger resistance values. In the meantime, the DC component of the signals remain constant for all resistors tested. **d**, Output voltage from the crystal oscillator showing the 32.768 kHz square wave. **e-g,** Voltage waveforms for different standard resistors after the HPF.

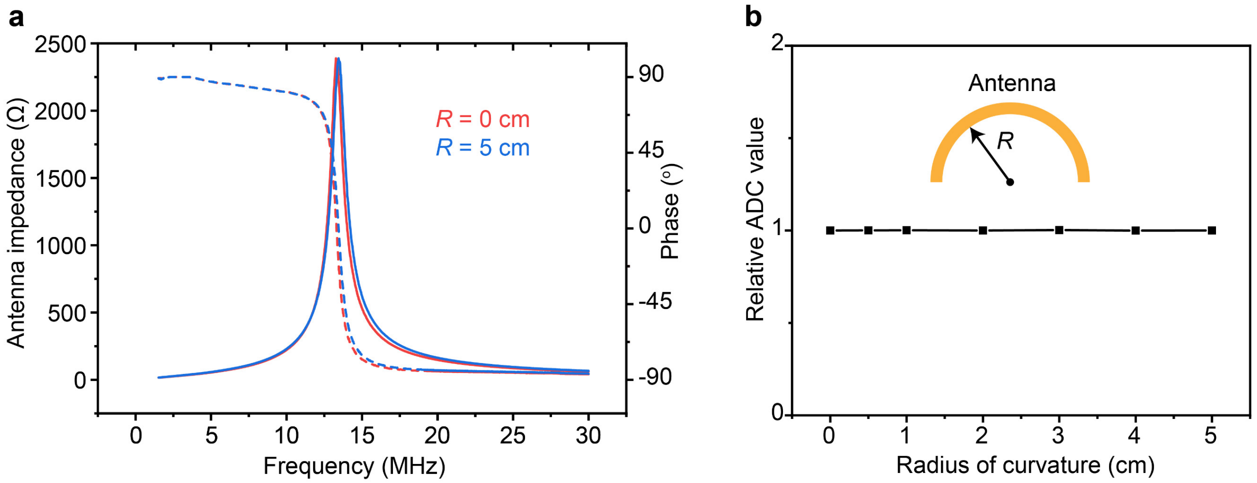

**Figure S3 | Antenna characterization under bending. a,** VNA scans of the same antenna with 0 cm and 5 cm bending radius. **b,** Wireless readout value of the ADC under different radius of curvature showing stable operature for the flexible antenna.

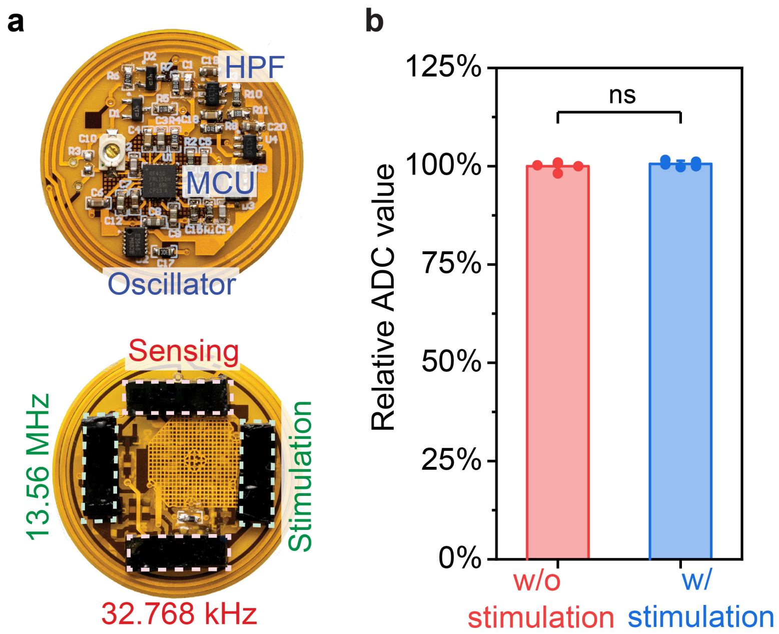

**Figure S4 | Validation of the sensing performance under simultaneous stimulation. a,** Photographs of the front (left) and back (right) sides of the smart bandage showing the microcontroller unit, crystal oscillator, high-pass filter, and stimulation and sensing electrodes. **b,** Relative ADC values of the 32.768 kHz impedance sensing channel with and without the 13.56 MHz stimulation electrodes connected.

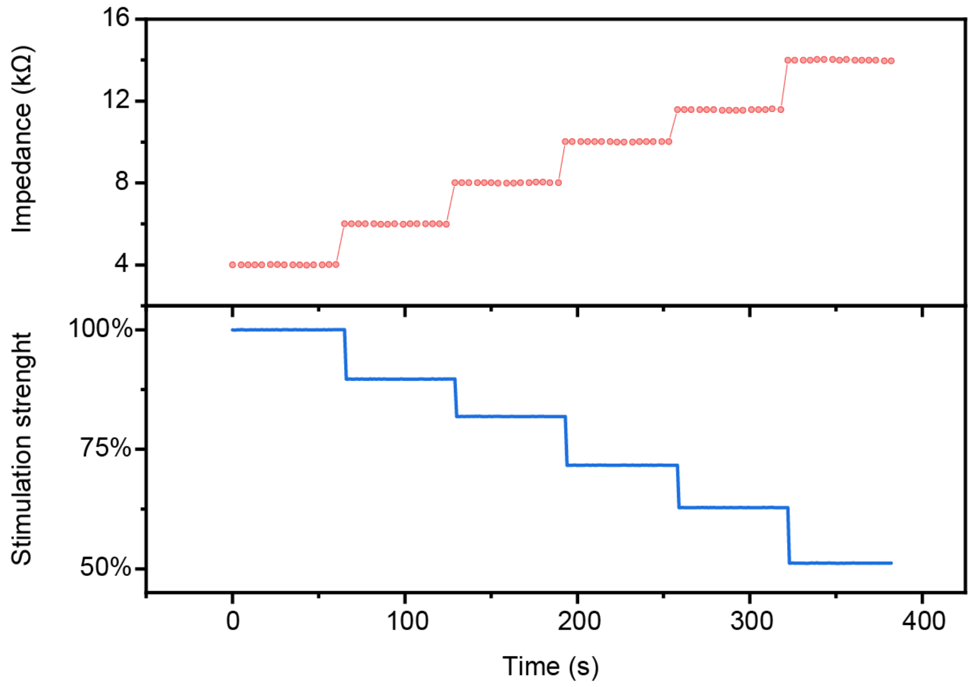

**Figure S5 | Closed-loop operation of the wireless smart bandage.** The reader can analyze the received sensing data (e.g., impedance) and adjust the emitted RF power in a closed-loop manner.

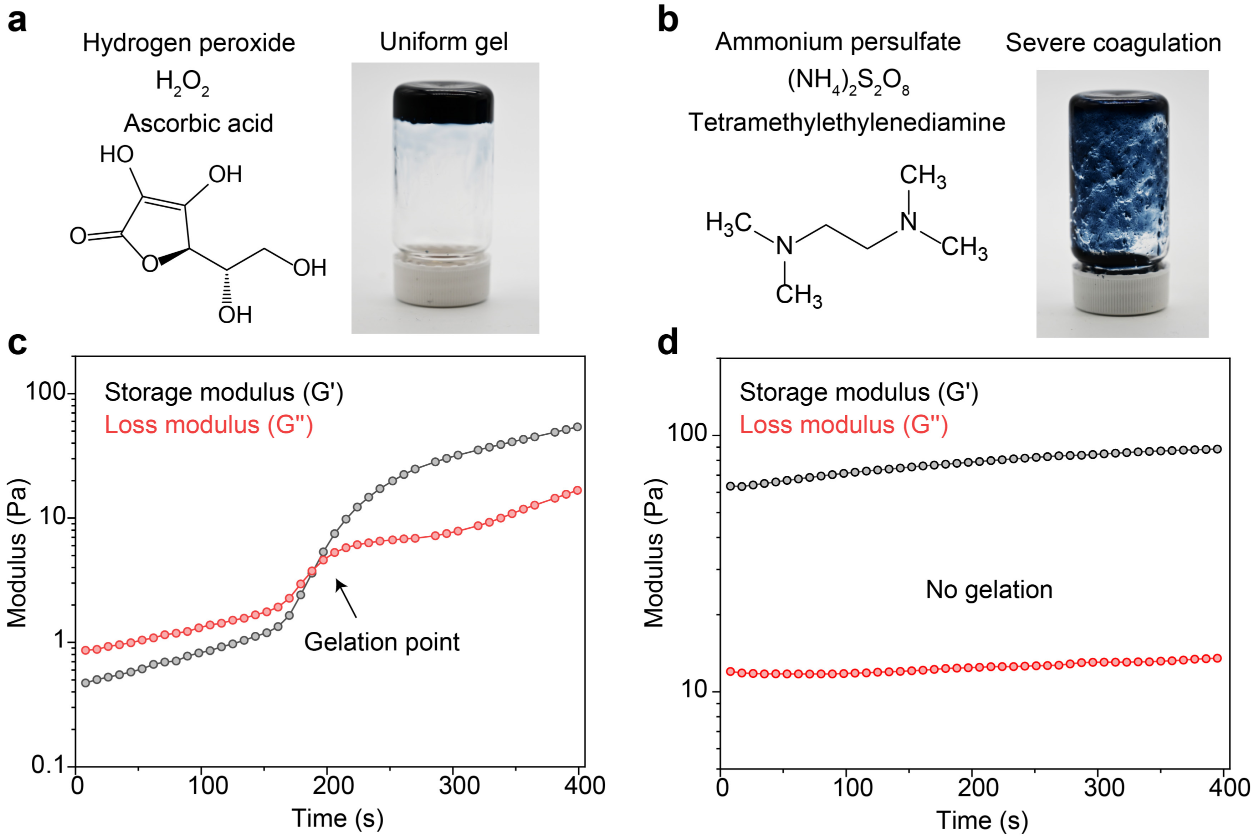

**Figure S6 | Comparison of the gelation outcome using different initiators. a** and **b**, chemical structures of different initator pairs (left) and corresponding photos (right) of the gels after reaction. **c** and **d**, Rheological measurements during the gelation process showing that only the non-ionic redox pair of hydrogen peroxide ($H_2O_2$) and ascorbic acid allows rapid and homogeneous gelation at room temperature (~3 min). Conventional radical initiators that contains both ionic and basic species, i.e., ammonium persulfate (AP) and *N,N,N′,N′*-tetramethylethylenediamine (TEMED) would cause severe coagulation of PEDOT:PSS.

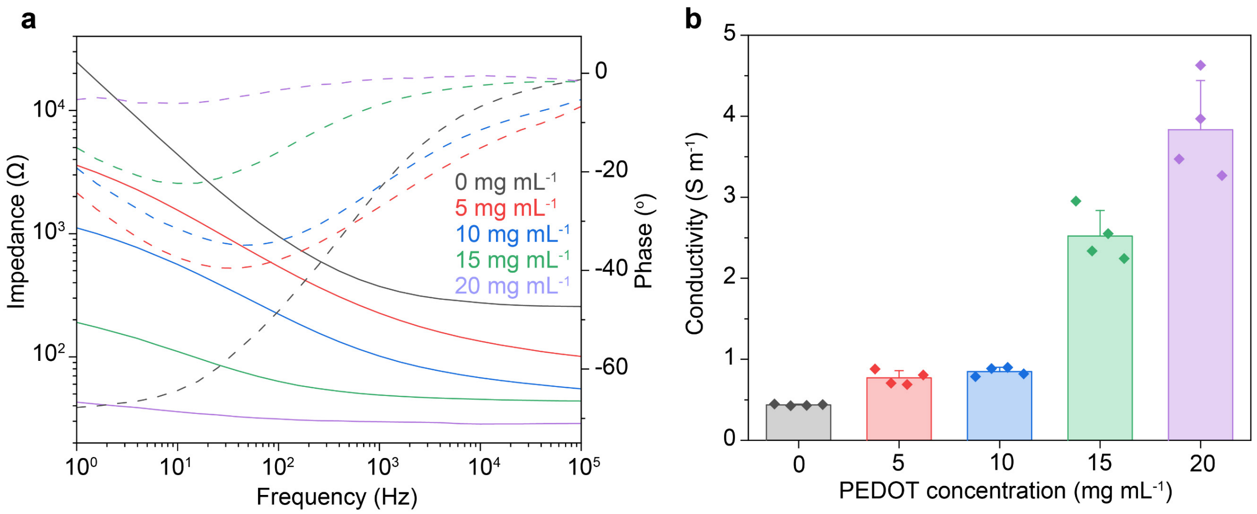

**Figure S7 | EIS measurement (a) and electrical conductivity (b) of the hydrogel electrode with different concentrations of PEDOT:PSS.** The hydrogel is consisting of 150 mg $mL^{-1}$ NIPAM, 12 mg $mL^{-1}$ AAm, and PEDOT:PSS from 0 to 20 mg $mL^{-1}$. In general, the higher the PEDOT:PSS concentration, the lower the impedance and the higher the conductivity. Additionally, the phase angle changed from a capacitive feature (i.e. high dependency of phase angle on frequency) for 0 mg $mL^{-1}$ of PEDOT:PSS to an almost resistive feature (i.e. phase angle independent of frequency) for 20 mg $mL^{-1}$ of PEDOT:PSS in the entire hydrogel. In all measurements, the contact hydrogel volume with PBS is $1\times1\times0.2$ $cm^3$, length×width×thickness.

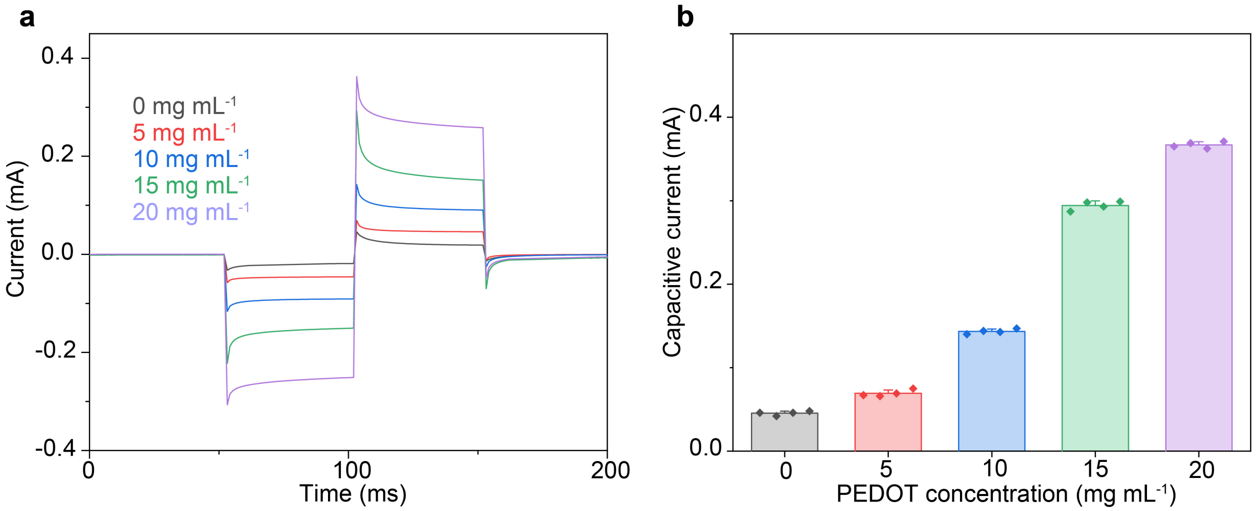

**Figure S8 | Chronoamperometry measurement (a) and injected current (b) of the hydrogel electrode with different concentrations of PEDOT:PSS.** The hydrogel is consisting of 150 mg $mL^{-1}$ NIPAM, 12 mg $mL^{-1}$ AAm, and PEDOT:PSS from 0 to 20 mg $mL^{-1}$. In general, the higher the PEDOT:PSS concentration, the better the charge injection as shown from the higher capacitive current determined by the initial spike at the current onset point. In all measurements, the contact hydrogel volume with PBS is 1×1×0.2 $cm^3$, length×width×thickness.

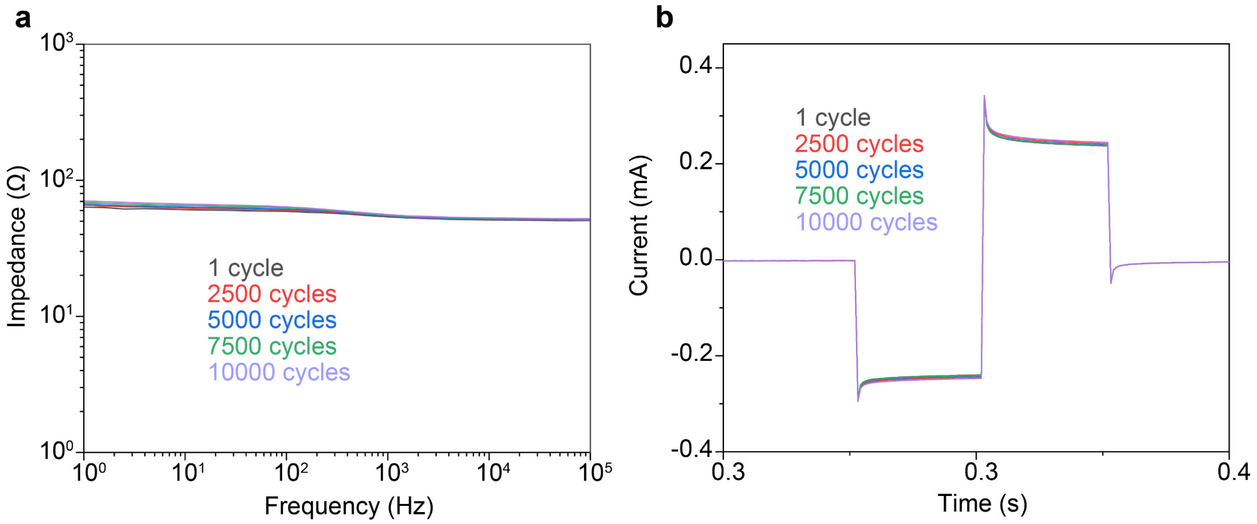

**Figure S9 | Cyclic tests of EIS (a) and chronoamperometry (b) of the hydrogel electrode showing the high electrochemical stability after at least 10000 cycles of repetitive charge injections.**

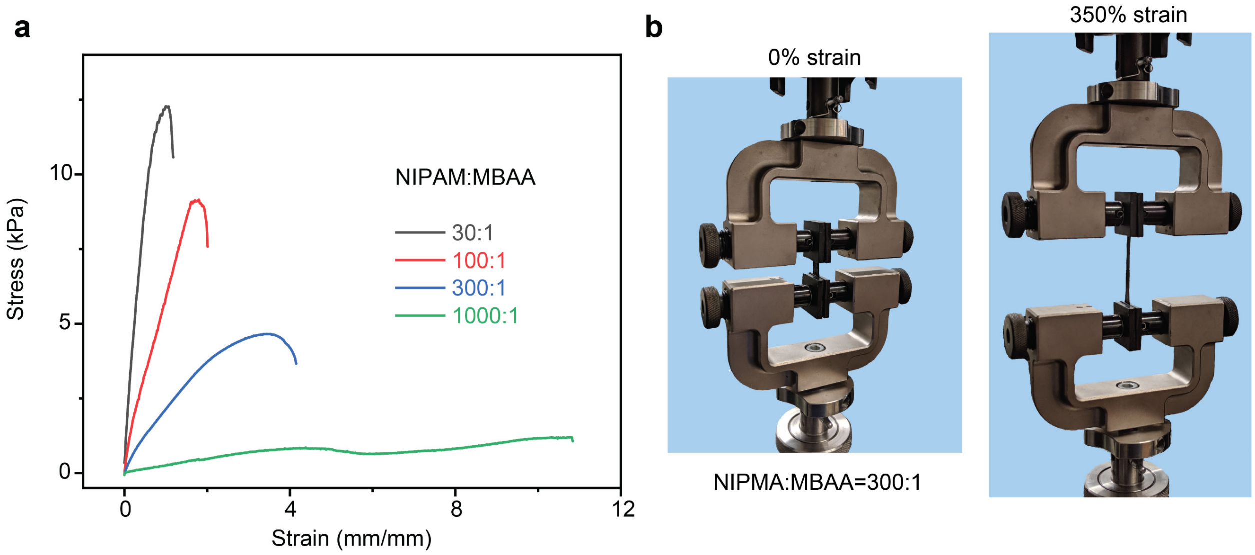

**Figure S10 | Uni-directional tensile test of the composite hydrogel electrode. a,** Stress-strain curves of composite hydrogels with different NIPAM over MBAA ratios, i.e., crosslinking densities. **b**, Photos showing the high stretchability of the hydrogel with the NIPAM over MBAA ratio at 300:1. The hydrogel is consisting of 150 mg $mL^{-1}$ NIPAM, 20 mg $mL^{-1}$ PEDOT:PSS, 12 mg $mL^{-1}$ AAm, and MBAA from 0.15 to 5 mg $mL^{-1}$.

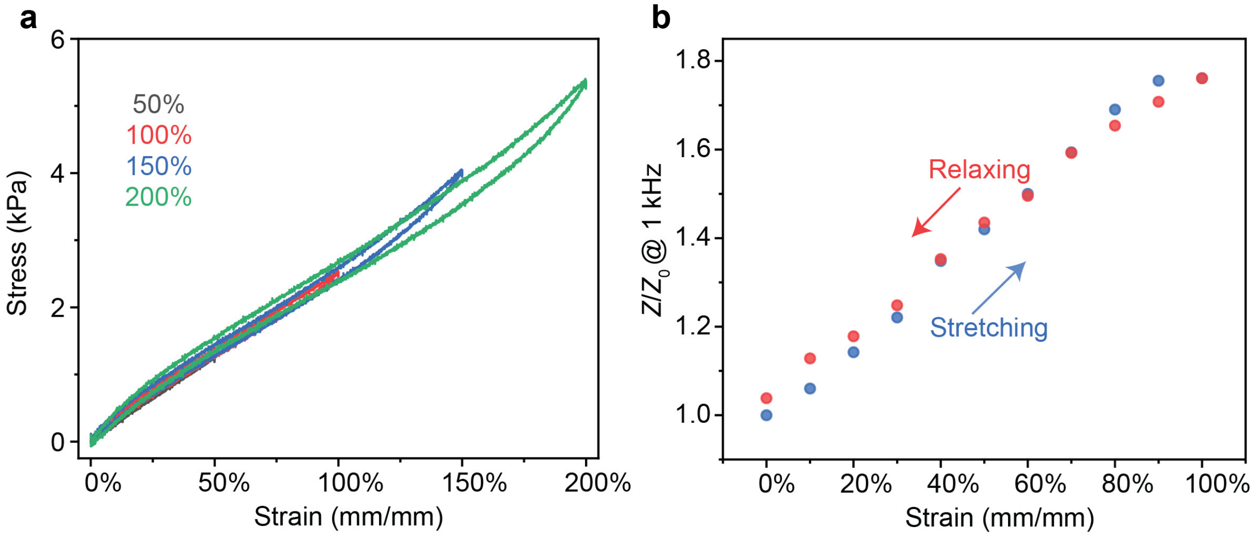

**Figure S11 | Cyclic loading of the hydrogel electrode and corresponding impedance changes.** **a**, Cyclic stress-strain curves of the same hydrogel being stretched to different strain levels and reversed. The covalently crosslinked poly(NIPAM-*ran*-AAm) network showed high elasticity with minimal hysteresis up to at least 200% strain. **b**, Impedance measurement of the same hydrogel being stretched to 100% strain and relaxed. The hydrogel is consisting of 150 mg $mL^{-1}$ NIPAM, 12 mg $mL^{-1}$ AAm, and 20 mg $mL^{-1}$ PEDOT:PSS.

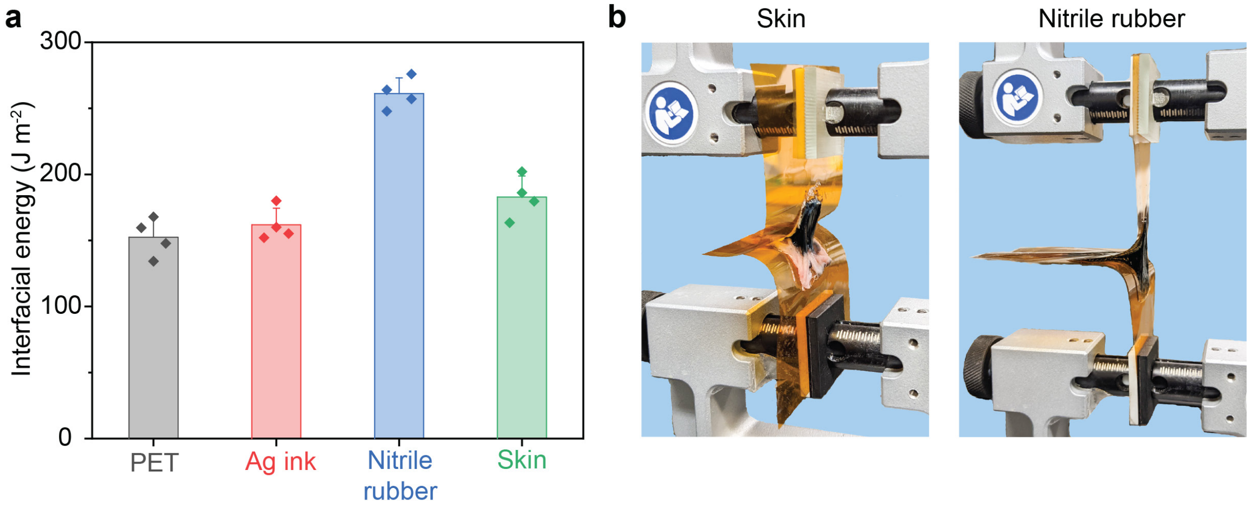

**Figure S12 | 180-degree peeling test of the hydrogel adhesion onto various substrates. a**, Quantitative comparison of the interfacial energy of the hydrogel electrode and different surfaces including plastic, metal, rubber, and tissue sample. **b**, Photos showing the tought adhesion of the hydrogel electrode on a mouse skin and nitrile rubber. The hydrogel is consisting of 150 mg $mL^{-1}$ NIPAM, 12 mg $mL^{-1}$ AAm, and 20 mg $mL^{-1}$ PEDOT:PSS. Ag ink was prepared by screen printing and drying. The skin tissue was harvested from mice.

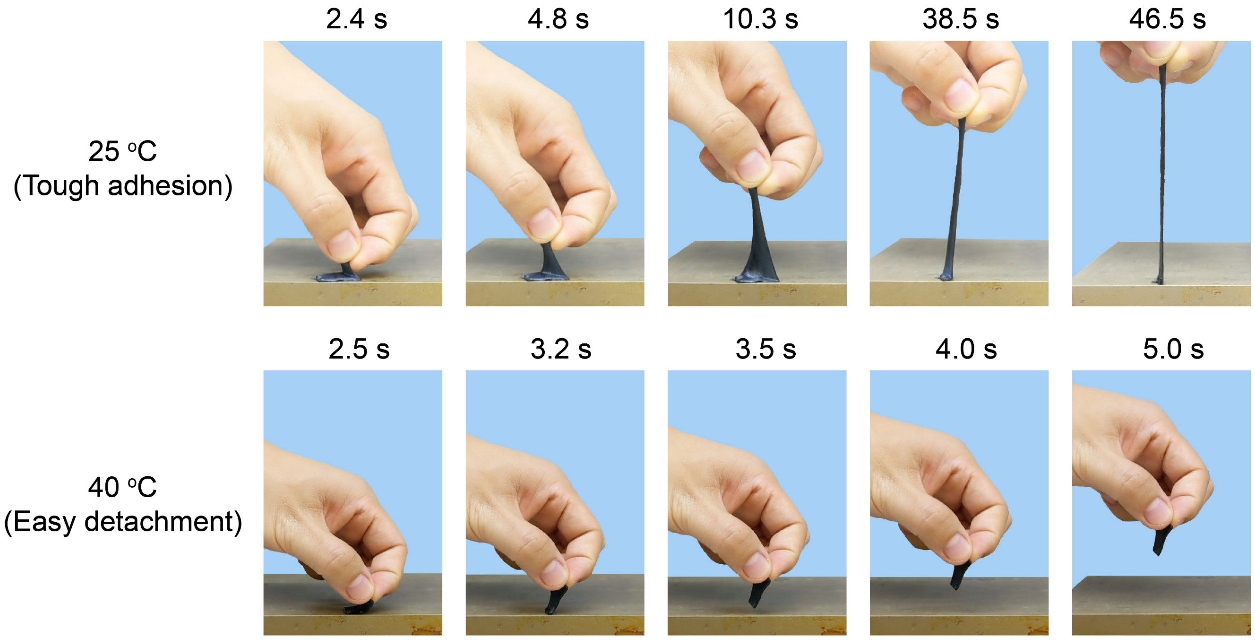

**Figure S13 | Photos showing the tunable adhesion of the same hydrogel electrode with a hot plate under different temperature.** The hydrogel exhibits tough adhesion at room temperature but completely loses its adhsion at 40 °C.

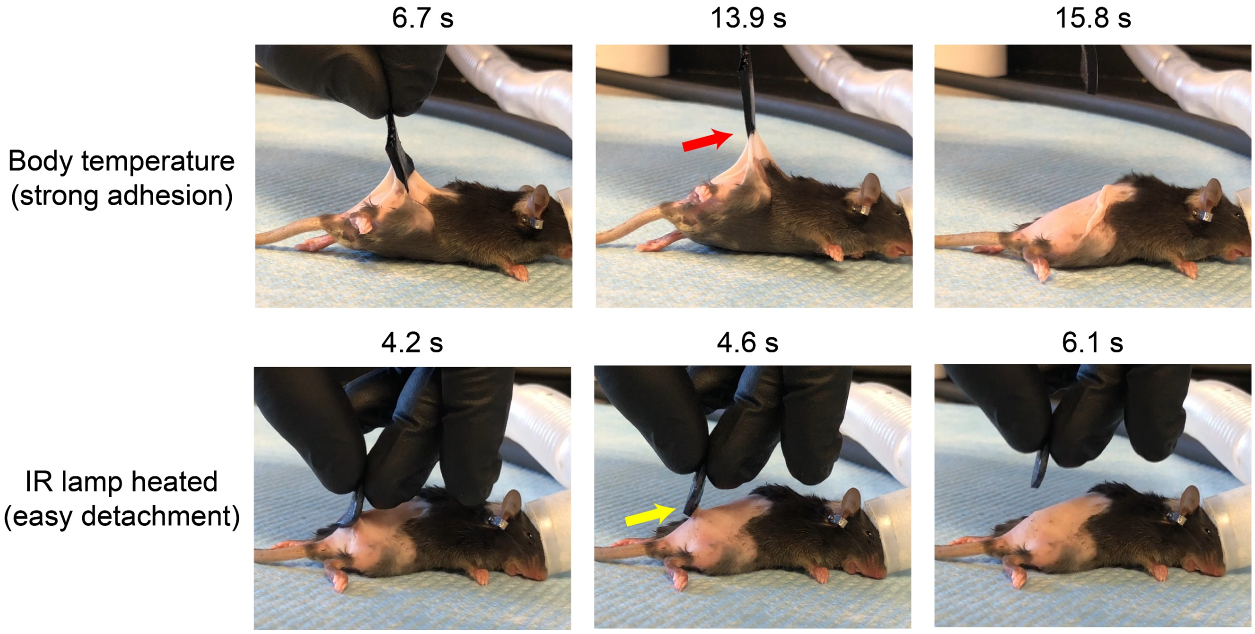

**Figure S14 | Photos showing the tunable adhesion of the same hydrogel electrode with moust skin under different temperature.** The hydrogel exhibits tough adhesion at room temperature but completely loses its adhsion after slight IR lamp heating.

| Host Response Score | 0 (none) | 1 (minimal) | 2 (mild) | 3 (moderate) | 4 (severe) |
|---|---|---|---|---|---|
| **Cell Response** | | | | | |
| Polymorpho-nuclear cells | 0 | 1-5/hpf | 5-10/hpf | Heavy infiltrate | Packed |
| Lymphocytes | 0 | 1-5/hpf | 5-10/hpf | Heavy infiltrate | Packed |
| Plasma cells | 0 | 1-5/hpf | 5-10/hpf | Heavy infiltrate | Packed |
| Macrophages | 0 | 1-5/hpf | 5-10/hpf | Heavy infiltrate | Packed |
| Multinucleated giant cells | 0 | 1-2/hpf | 3-5/hpf | Heavy infiltrate | Sheets |
| Necrosis | 0 | Minimal | Mild | Moderate | Severe |
| **Tissue Response** | | | | | |
| Neo-vascularization | 0 | Minimal capillary proliferation, focal, 1-3 buds | Groups of 4-7 capillaries with supporting fibroblastic structures | Broad band capillaries with supporting fibroblastic structures | Extensive band of capillaries with supporting fibroblastic structures |
| Fibrosis | 0 | Narrow band | Moderately thick band | Thick band | Extensive band |
| Fatty infiltration | 0 | Minimal amount of fat associated with fibrosis | Several layers of fat and fibrosis | Elongated and broad accumulation of fat cells about the implant site | Extensive fat completely surrounding the implant |

**Table S2 | Semi-quantitative scoring criteria for paraffin sections to assess the hydrogel biocompatibility.** hpf stands for high-powered field, i.e., 40× objective.

**a**

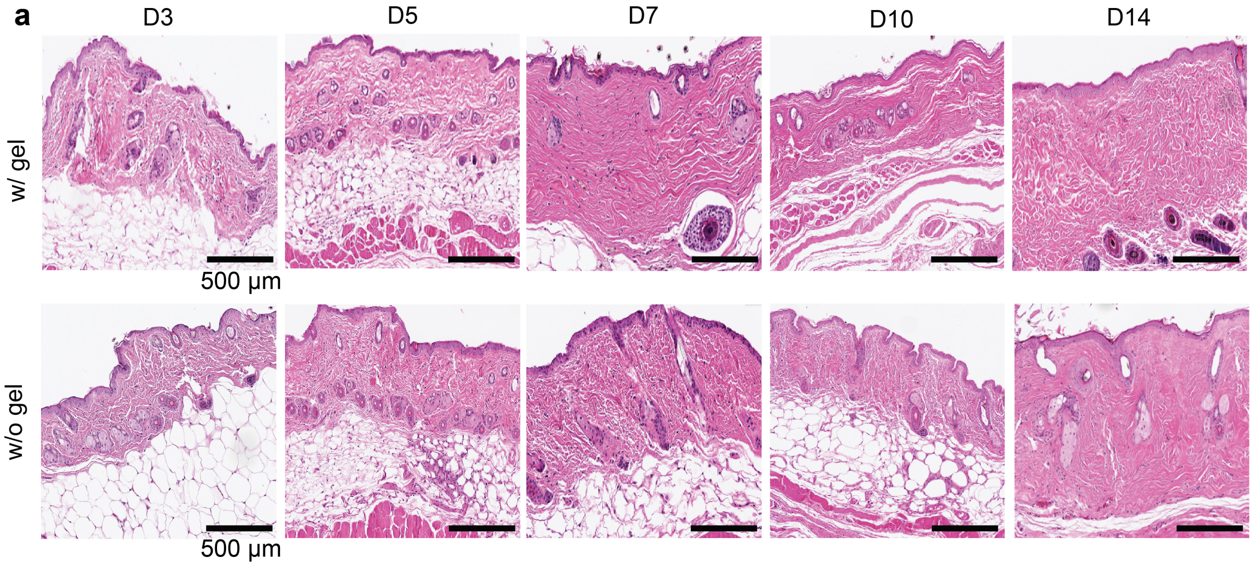

**b**

| Time | Day 3 | | Day 5 | | Day 7 | | Day 10 | | Day 14 | |
|---|---|---|---|---|---|---|---|---|---|---|
| Animal group | w/o gel | w/ gel | w/o gel | w/ gel | w/o gel | w/ gel | w/o gel | w/ gel | w/o gel | w/ gel |
| Polymorphonuclear cells | 0 | 0 | 0 | 0 | 0 | 0 | 0 | 0 | 0 | 0 |
| Lymphocytes | 1 | 1 | 1 | 0.67 | 1 | 0.67 | 1 | 1 | 1 | 1 |
| Plasma cells | 0 | 0 | 0 | 0.67 | 0 | 0 | 0 | 0 | 0.33 | 0 |
| Macrophages | 0.5 | 0.5 | 0.33 | 0.33 | 0.33 | 0 | 0 | 0 | 0.33 | 0 |
| Giant cells | 0 | 0 | 0 | 0 | 0 | 0 | 0 | 0 | 0 | 0 |
| Necrosis | 0 | 0 | 0 | 0 | 0 | 0 | 0 | 0 | 0 | 0 |
| Subtotal A (x2) | 3 | 3 | 1.33 | 1.67 | 1.33 | 0.67 | 1 | 1 | 1.67 | 1 |
| Fibrosis | 0 | 0.5 | 0.67 | 0 | 0 | 0 | 0 | 0 | 0 | 0 |
| Subtotal B | 0 | 0.5 | 2 | 0 | 0 | 0 | 0 | 0 | 0 | 0 |
| Total (Subtotal A + Subtotal B) | 3 | 3.5 | 2 | 1.67 | 1.33 | 0.67 | 1 | 1 | 1.67 | 1 |

**c**

| Reactivity Calculation | | | |
|---|---|---|---|
| | w/o gel | w/ gel | Relative Score (w/ gel – w/o gel) |
| Day 3 | 3 | 3.5 | 0.5 |
| Day 5 | 2 | 1.67 | 0 |
| Day 7 | 1.33 | 0.67 | 0 |
| Day 10 | 1 | 1 | 0 |
| Day 14 | 1.67 | 1 | 0 |

| Reactivity Grade | Test Sample Relative Score |
|---|---|
| Minimal or no reaction | 0.0 – 2.9 |
| Slight reaction | 3.0 – 8.9 |
| Moderate reaction | 9.0 – 15.0 |
| Severe reaction | > 15 |

**Figure S15 | Hydrogel electrode showing excellent biocompatibility after long-term contact with mouse skin. a,** H&E cross-sections of mouse skin tissue with and without hydrogels over the course of two weeks showing no visible inflammatory response. **b,** Average immune cell counts over time showing minimal or no reaction quantitatively. **c,** Scoring of the immune response reactivity showing minimal/no reaction for the hydrogel electrode. The hydrogel is consisting of 150 mg mL$^{-1}$ NIPAM, 12 mg mL$^{-1}$ AAm, and 20 mg mL$^{-1}$ PEDOT:PSS.

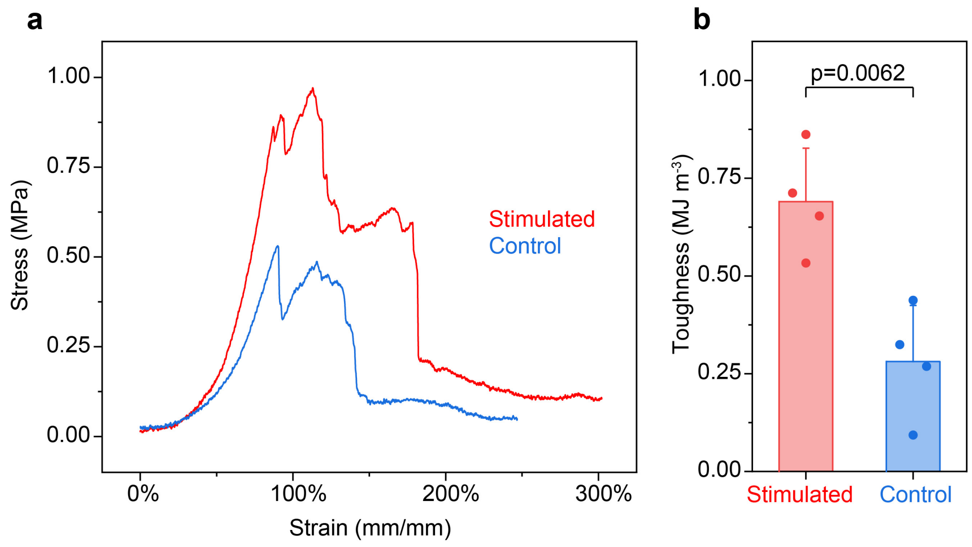

**Figure S16 | Electrical stimulation induces higher skin toughness after healing compared to the control tissue.** Tensile test (**a**) and corresponding toughness (**b**) of healed skin samples samples after 13 days with and without electrical stimulation. Toughness values were calculated by integrating the underlying area of the stress-strain curves.

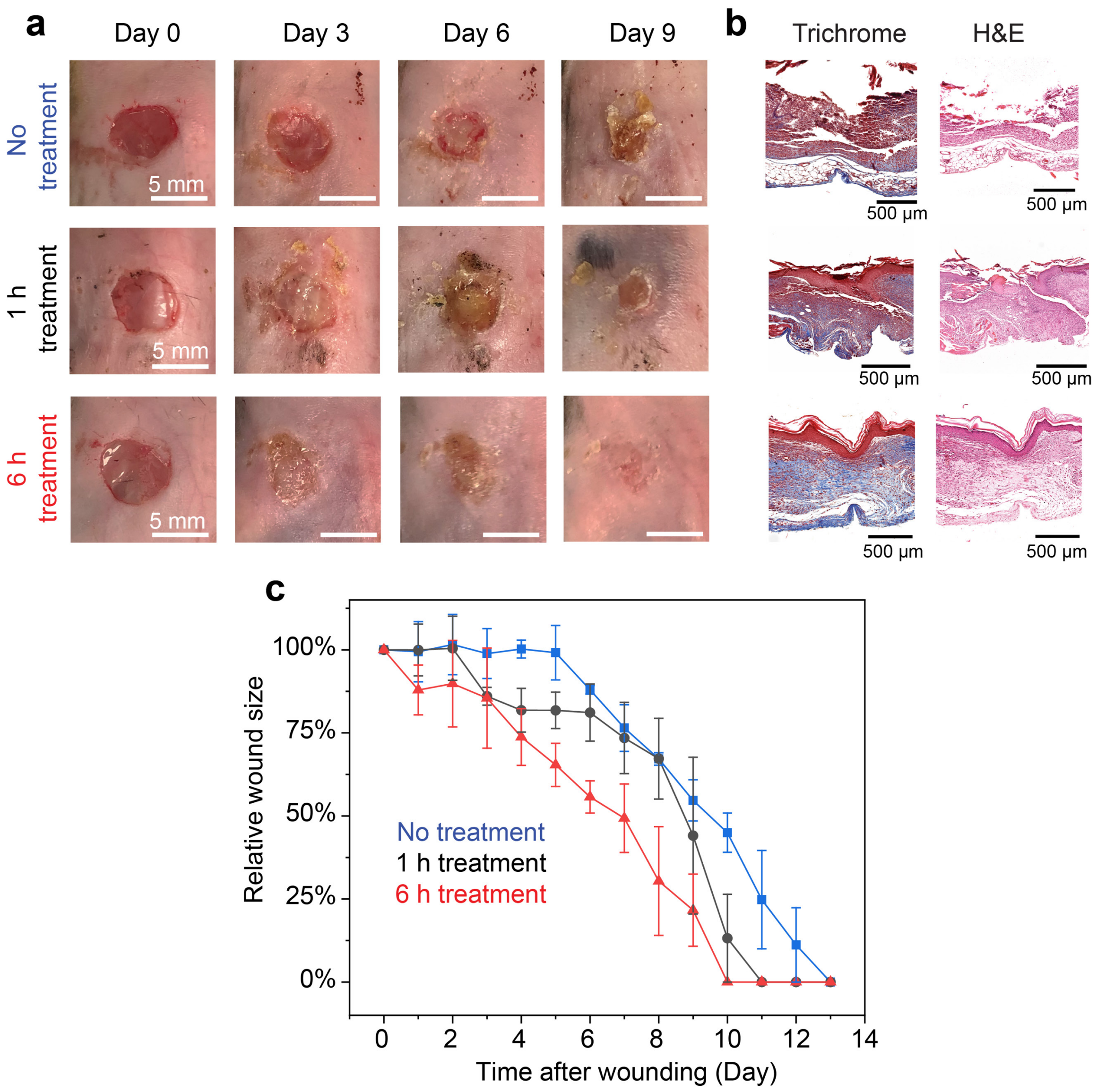

**Figure S17 | Wireless smart bandage allows longer treatment per day with better healing outcomes. a,** Representative photos showing the progression of excisional wounds with different treatment durations. **b,** Representative cross-sectional histology images of skin tissue harvested from mice with different treatment periods. Left, Masson's trichrome, where a visible intact epidermal and dermal layer observed in stimulated treatment group in the Masson's trichrome stain, visualized by a red surface layer which stains for muscle cells and blue layer below, which stains for collagen; Right, hematoxylin and eosin (H&E). **c,** Relative wound size curve, indicating accelerated tissue regeneration with logner treatment duration. $n$=5 for each group. All data are represented as mean ± standard deviation.

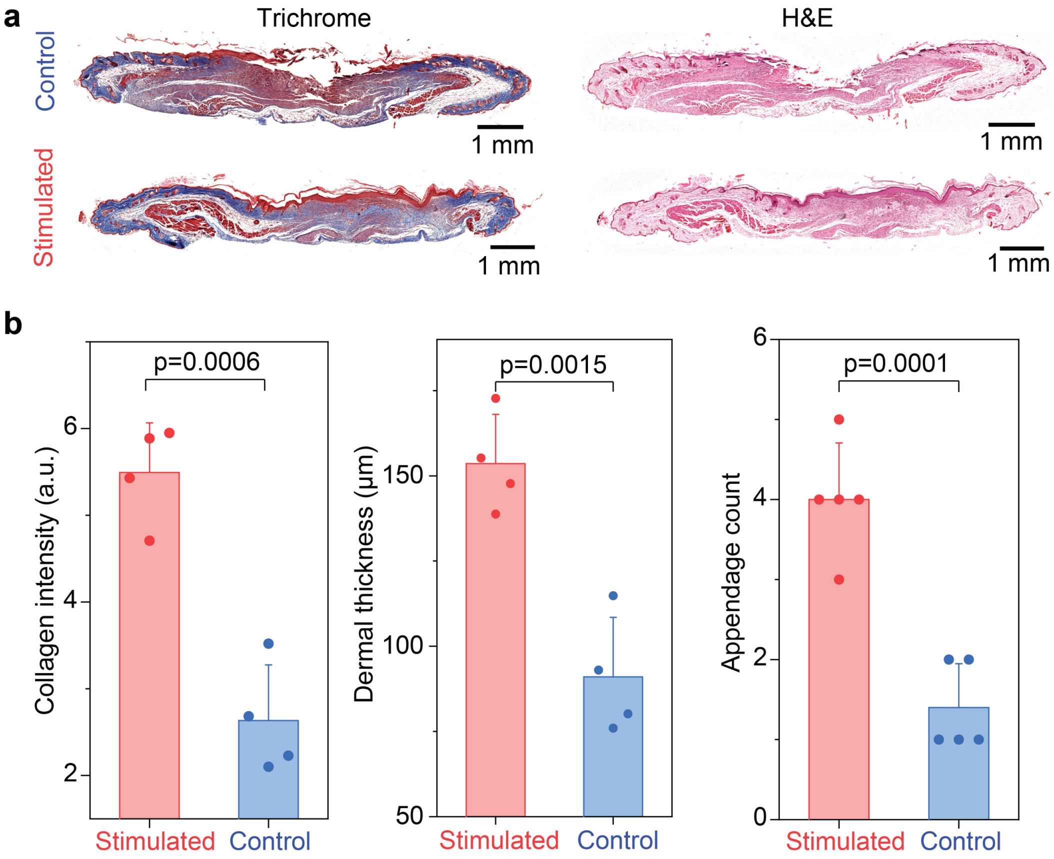

**Figure S18 | Histological studies show improved wound healing outcomes for excisional wounds treated with electrical stimulation. a,** Representative cross-sectional histology images of skin tissues harvested from mice with and without electrical stimulation after 13 days. A visible intact epidermal and dermal layer observed in stimulated treatment group in the Masson's trichrome stain, visualized by a red surface layer which stains for muscle cells and blue layer below, which stains for collagen. $n$=4 for each group. **b,** Quantitative evaluations showing increased collagen intensity, dermal thickness, and appendage count for the stimulated group. All data are represented as mean ± standard deviation. Two-tailed t-test assuming equal variances were performed for the $p$ values.

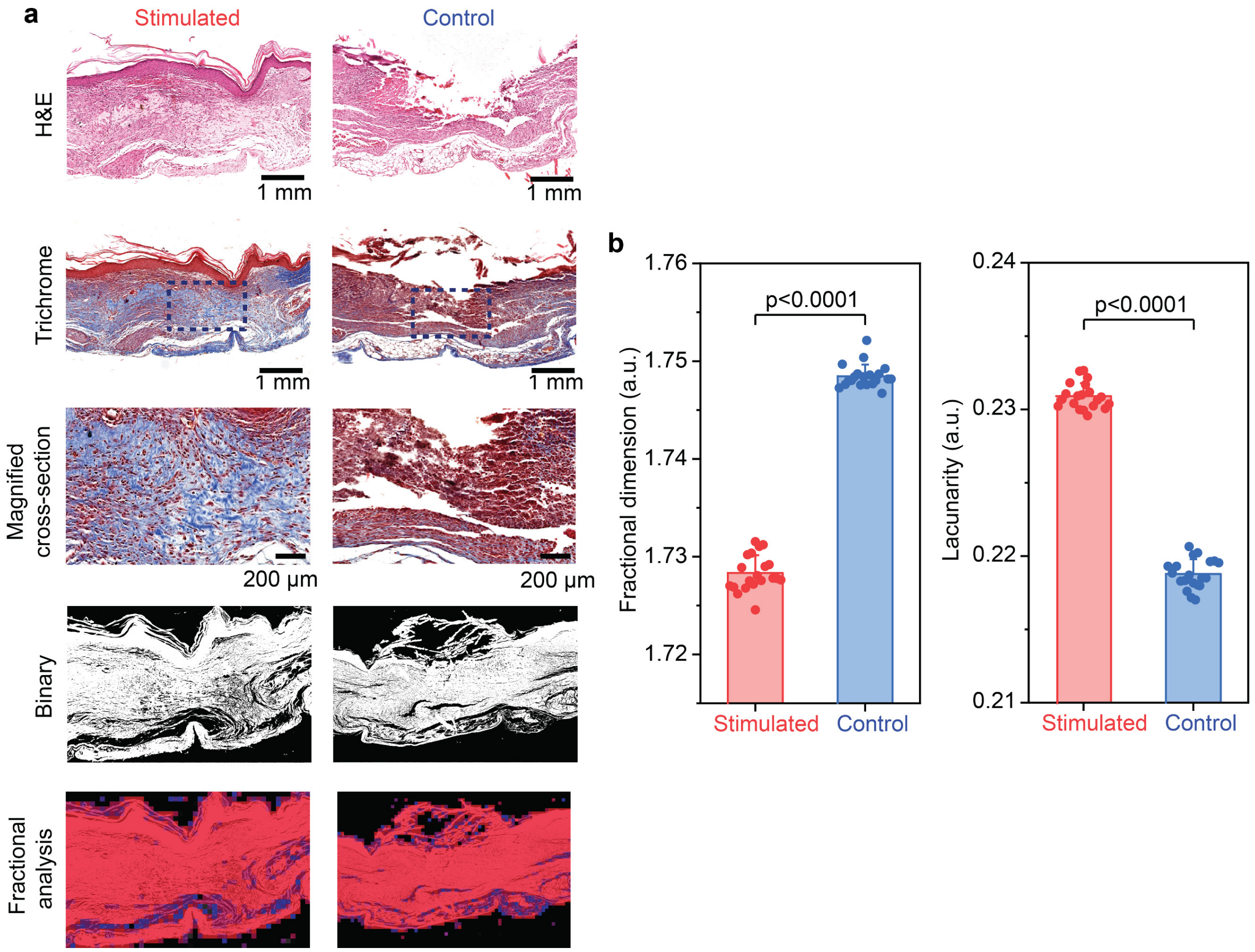

**Figure S19 | FracLac evaluation shows less scar-like fiber arrangement for excisional wounds treated with electrical stimulation after 13 days. a,** Representative cross-sectional histology images and corresponding FracLac of skin tissues harvested from mice with and without electrical stimulation. Magnified cross-section, outlined with a dotted rectangle, highlights the collagen composition and microvessel formation of healed tissue. Masson's trichrome images were binarized using FracLac and collagen denisty was quantified, represented by red and blue colored boxes in the bottom-most row. **b,** Quantitative comparison of local fractal dimension (FD) and lacunarity. FD measures density of collagen networks. A higher FD has a denser and scar-like fiber arrangement. Lacunarity measures the amount of randomness or heterogeneity in a sample. A low lacunarity implies less heterogenous collagen fiber orientation. All data are represented as mean ± standard deviation. Two-tailed t-test assuming equal variances were performed for the *p* values.

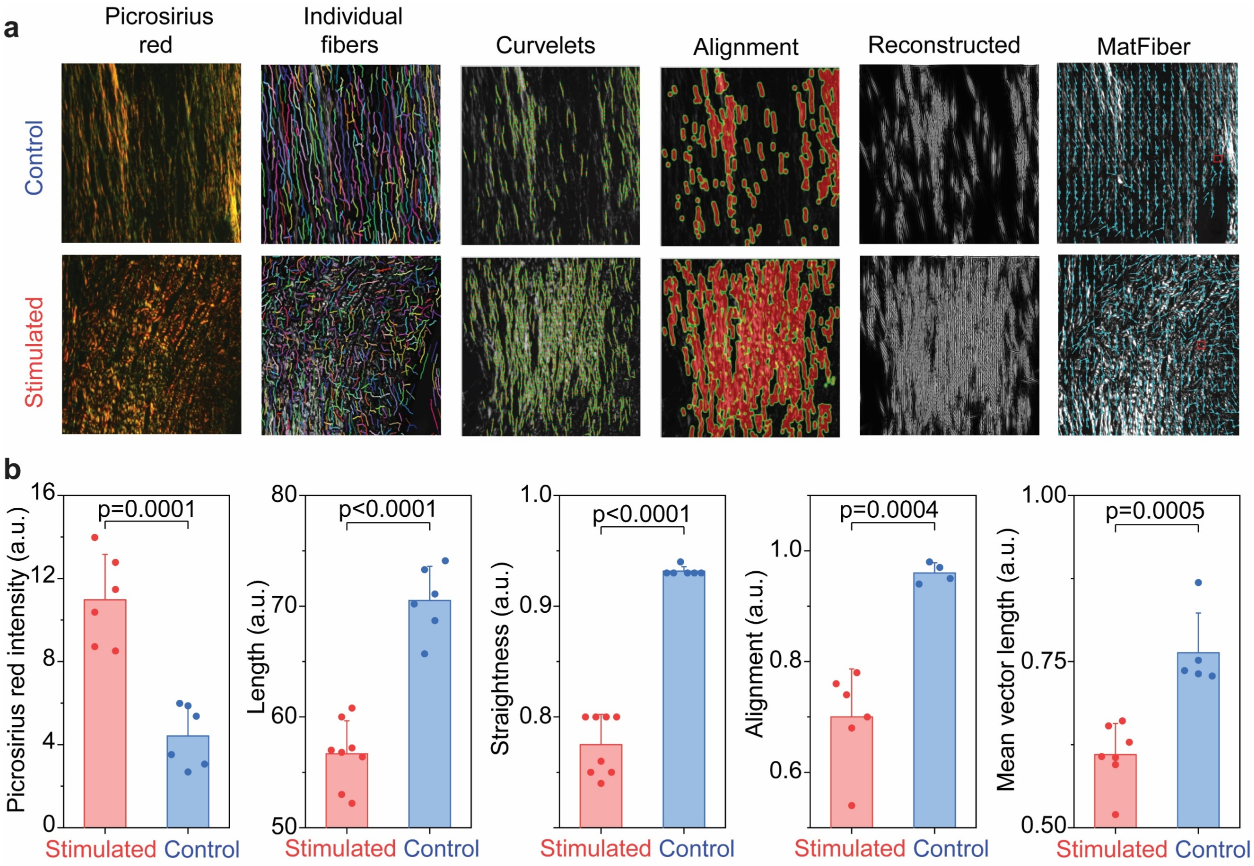

**Figure S20 | Picrosirius red staining shows more random collagen networks for excisional wounds treated with electrical stimulation after 13 days. a,** Representative picrosirius red stained images and collagen network analysis of skin tissues harvested from mice with and without electrical stimulation using Curve Align, CT-Fire and MatFiber. CT-FIRE algorithm outlines individual fibers to evaluate length, width, angle, and curvature. CurveAlign quantifies all fiber angles and strength of alignment within an image, tracing the fibers in green quantifying the fiber density with red heat maps and reconstructing the image in binary form to produce an output. MatFiber quantifies fiber alignment, tracing the fibers with blue arrows, reported as mean vector length. The strength of alignment ranges from a value of 0 (completely random fiber alignment) to 1 (completely aligned fibers). **b,** Quantitative comparison of picrosirius red intensity, collagen length, collagen straightness, collagen alignment, and mean vector length shows more random collagen networks and less fibrotic tissue formation. All data are represented as mean ± standard deviation. Two-tailed t-test assuming equal variances were performed for the *p* values.

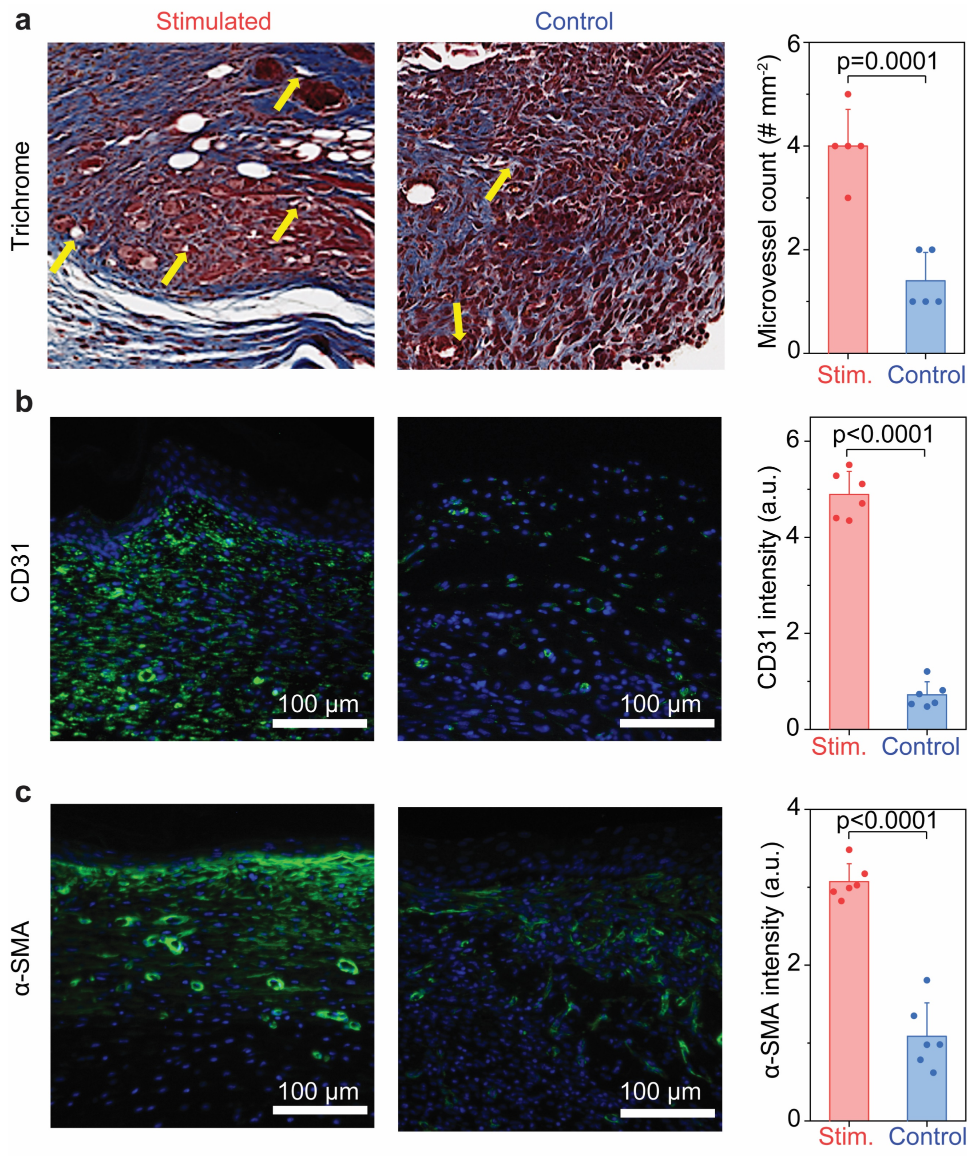

**Figure S21 | Histology and immunostaining reveals more neovascularization for excisional wounds treated with electrical stimulation after 13 days.** Representative Masson's trichrome (**a**), stained images and corresponding analysis showing microvessel count (indicated with yellow arrows), and vascularity of skin tissues harvested from mice with and without electrical stimulation stained with (**b**), CD31, which stains for blood vessels and (**c**), α-SMA, a marker for smooth muscle cells, namely myofibroblasts, which are commonly found in the walls of blood vessels. All data are represented as mean ± standard deviation. Two-tailed t-test assuming equal variances were performed for the *p* values.

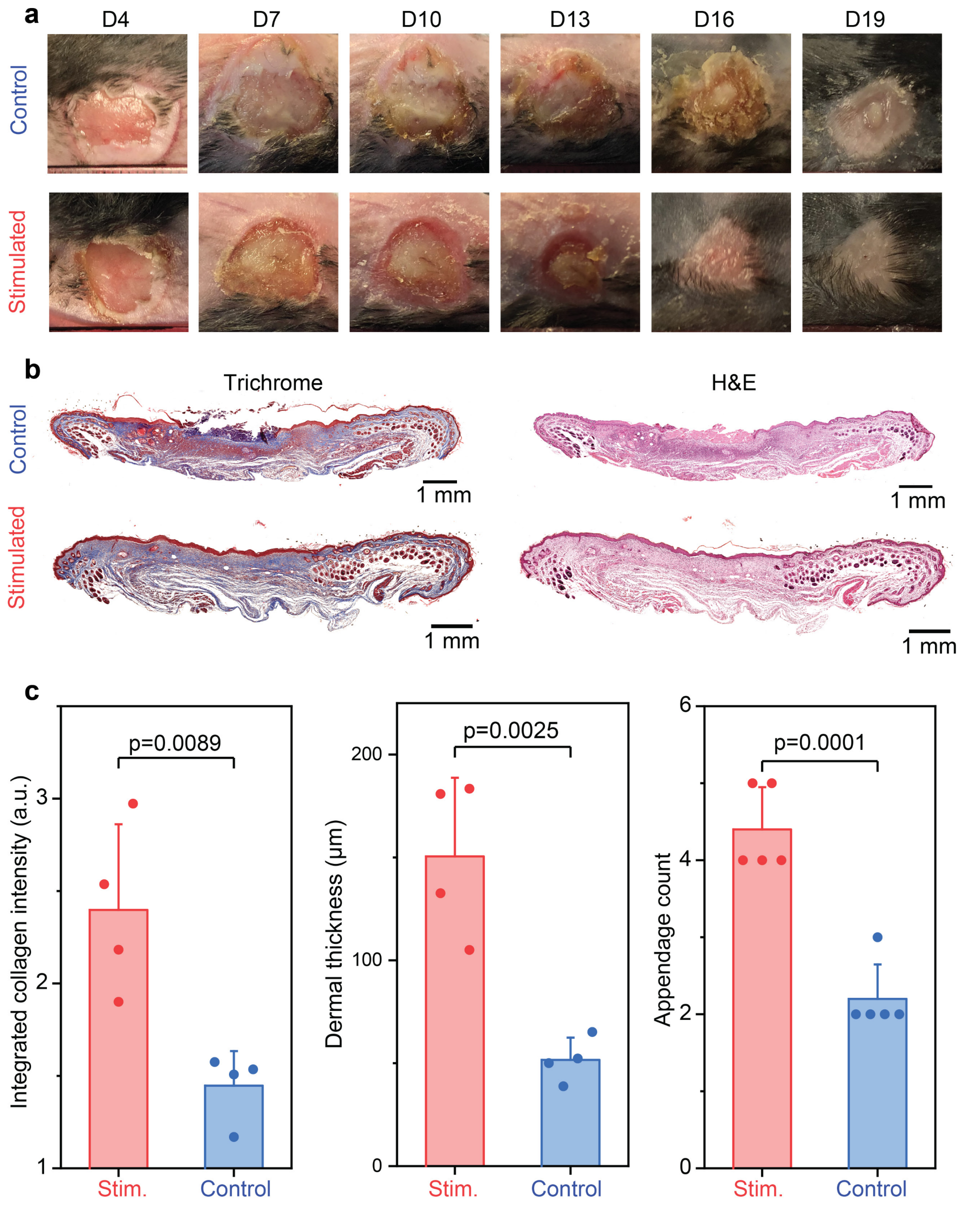

**Figure S22 | Histological studies show improved healing outcomes for burn wounds treated with electrical stimulation after 19 days. a,** Representative photos showing the progression of burn wounds with and without electrical stimulation. Stimulated wounds show visual signs of reduced necrotic tissue, observed with lighter-colored tissue surrounding the wound. Stimulated wounds also have accelerated dermal recorvery, as indicated by earlier scab formation (Day 13) and re-epithelialization of the wound tissue (Day 16). $n$=4 for each group **b,** Representative cross-sectional histology images of skin tissues harvested from mice with and without electrical stimulation. A visible intact epidermal and dermal layer observed in stimulated treatment group in the Masson's trichrome stain, visualized by a red surface layer which stains for muscle cells and blue layer below, which stains for collagen. **c,** Quantitative comparison of the histology images shows increased collagen intensity, dermal thickness, and appendage count for the stimulated group. All data are represented as mean ± standard deviation. Two-tailed t-test assuming equal variances were performed for the $p$ values.

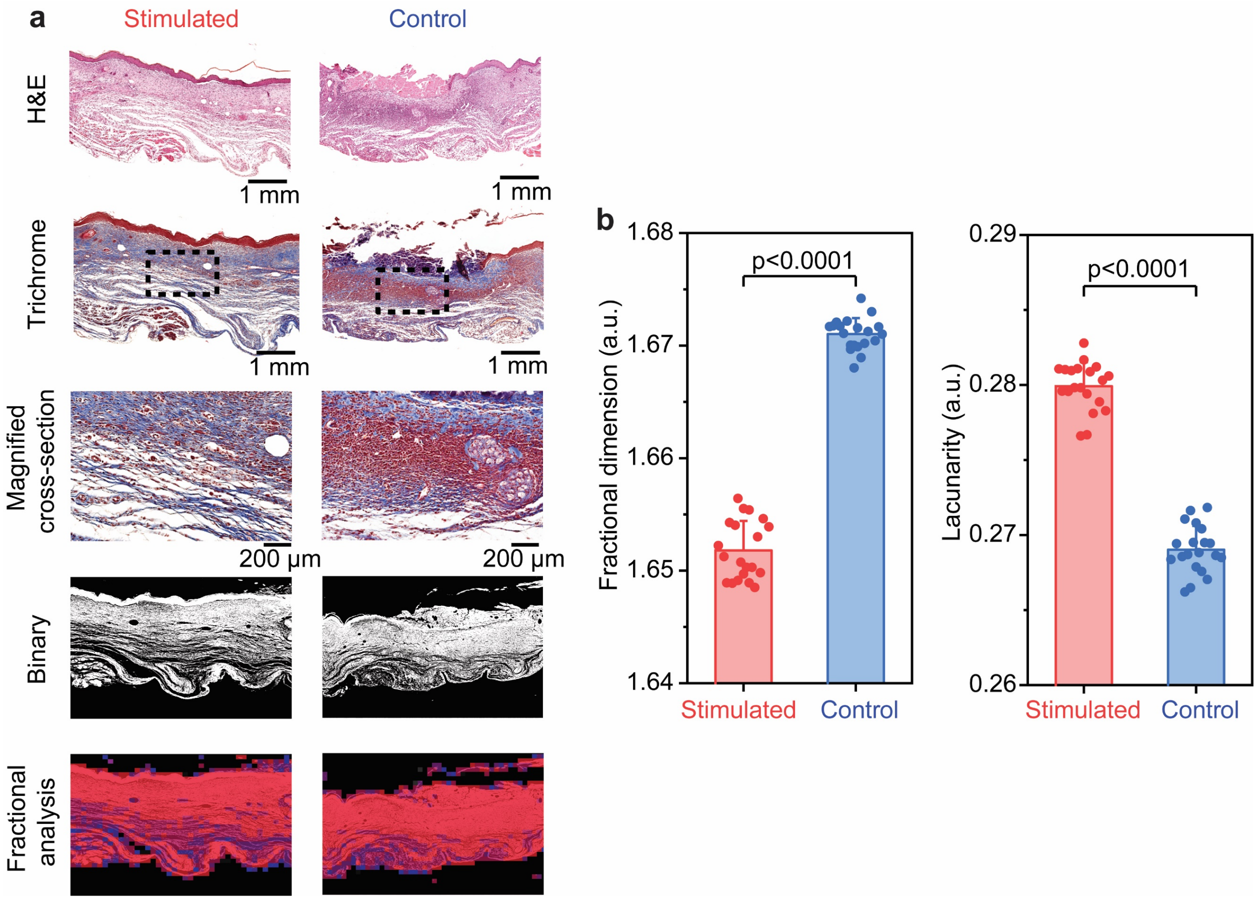

**Figure S23 | FracLac evaluation shows less scar-like fiber arrangement for burn wounds treated with electrical stimulation after 19 days. a,** Representative cross-sectional histology images and corresponding FracLac of skin tissues harvested from mice with and without electrical stimulation. Magnified cross-section, outlined with a dotted rectangle, highlights the collagen composition and microvessel formation of healed tissue. Masson's trichrome images were binarized using FracLac and collagen denisty was quantified, represented by red and blue colored boxes in the bottom-most row. **b,** Quantitative comparison of local fractal dimension (FD) and lacunarity. FD measures density of collagen networks. A higher FD has a denser and scar-like fiber arrangement. Lacunarity measures the amount of randomness or heterogeneity in a sample. A low lacunarity implies less heterogenous collagen fiber orientation. All data are represented as mean ± standard deviation. Two-tailed t-test assuming equal variances were performed for the *p* values.

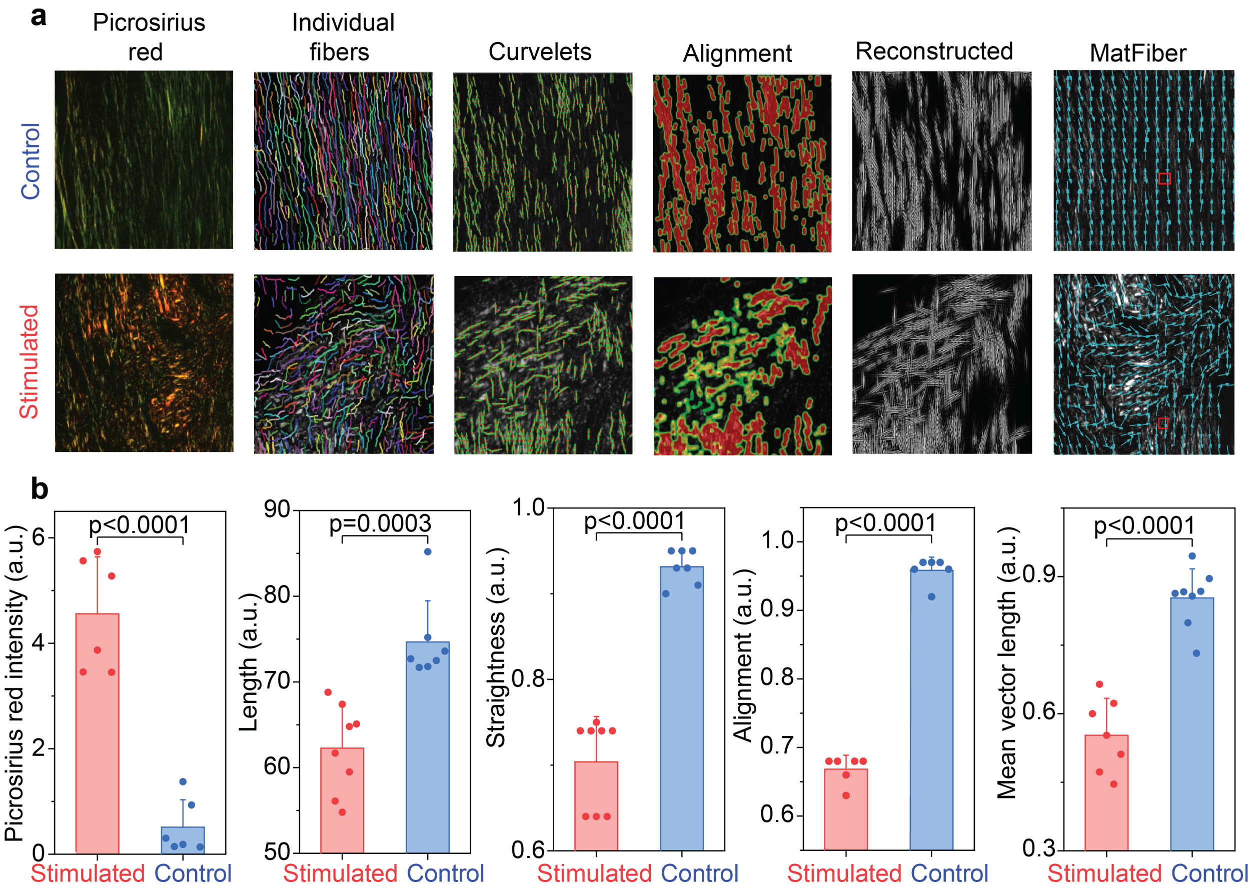

**Figure S24 | Picrosirius red staining shows more random collagen networks for burn wounds treated with electrical stimulation after 19 days. a,** Representative picrosirius red stained images and collagen network analysis of skin tissues harvested from mice with and without electrical stimulation using Curve Align, CT-Fire and MatFiber. CT-FIRE algorithm outlines individual fibers to evaluate length, width, angle, and curvature. CurveAlign quantifies all fiber angles and strength of alignment within an image, tracing the fibers in green quantifying the fiber density with red heat maps and reconstructing the image in binary form to produce an output. MatFiber quantifies fiber alignment, tracing the fibers with blue arrows, reported as mean vector length. The strength of alignment ranges from a value of 0 (completely random fiber alignment) to 1 (completely aligned fibers). **b,** Quantitative comparison of picrosirius red intensity, collagen length, collagen straightness, collagen alignment, and mean vector length shows more random collagen networks and less fibrotic tissue formation. All data are represented as mean ± standard deviation. Two-tailed t-test assuming equal variances were performed for the *p* values.

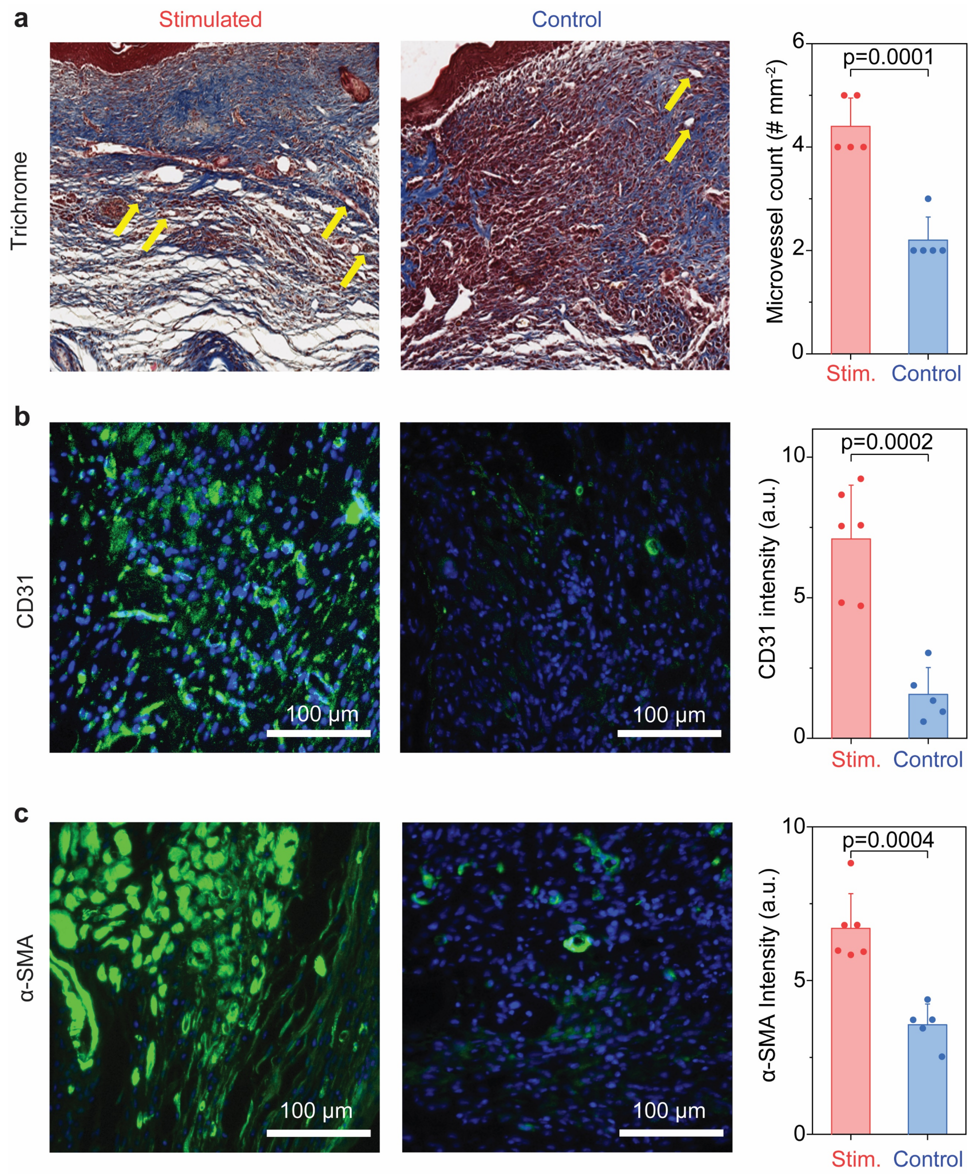

**Figure S25 | Histology and immunostaining reveals more neovascularization for burn wounds treated with electrical stimulation after 19 days.** Representative Masson's trichrome (**a**), stained images and corresponding analysis showing microvessel count (indicated with yellow arrows), and vascularity of skin tissues harvested from mice with and without electrical stimulation stained with (**b**), CD31, which stains for blood vessels and (**c**), α-SMA, a marker for smooth muscle cells, namely myofibroblasts, which are commonly found in the walls of blood vessels. All data are represented as mean ± standard deviation. Two-tailed t-test assuming equal variances were performed for the *p* values.

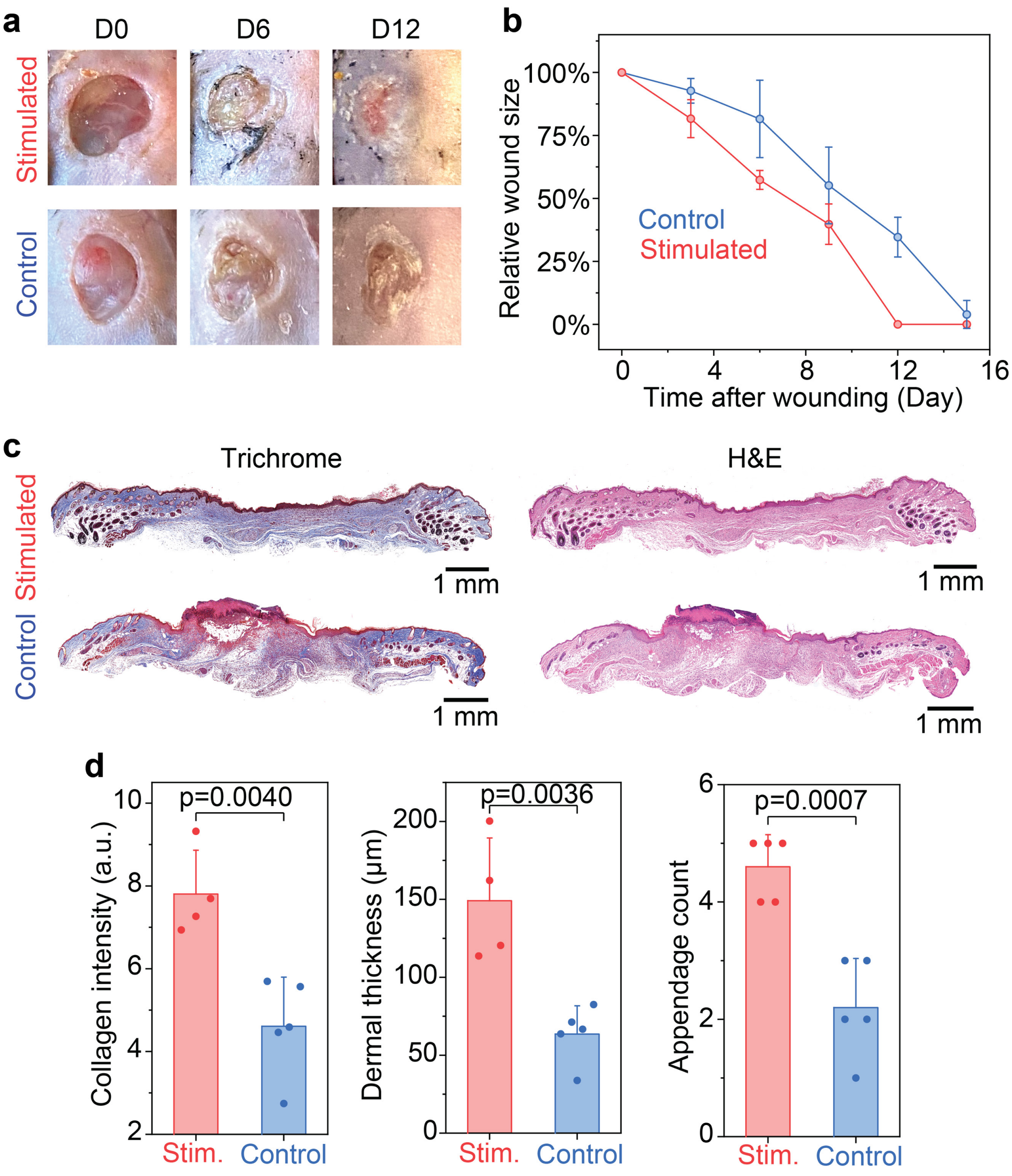

**Figure S26 | Histological studies show improved healing outcomes for STZ-induced diabetic excisional wound model treated with electrical stimulation after 15 days. a,** Representative photos showing the progression of excistional wounds in STZ-induced diabetic mice with and without electrical stimulation. Stimulated wounds show accelerated re-epitheliazation, visually observed with the formation of a scab (Day 6) and accelrated re-epitheliazation (Day 12). **b,** Relative size of the wound over time, indicating accelerated tissue regeneration with stimulation. $n$=4 for each group. All data are represented as mean ± standard deviation. **c,** Representative cross-sectional histology images of skin tissues harvested from mice with and without electrical stimulation. **d,** Quantitative comparison of the histology images showing increased collagen intensity, dermal thickness, and appendage count for the stimulated group. All data are represented as mean ± standard deviation. Two-tailed t-test assuming equal variances were performed for the $p$ values.

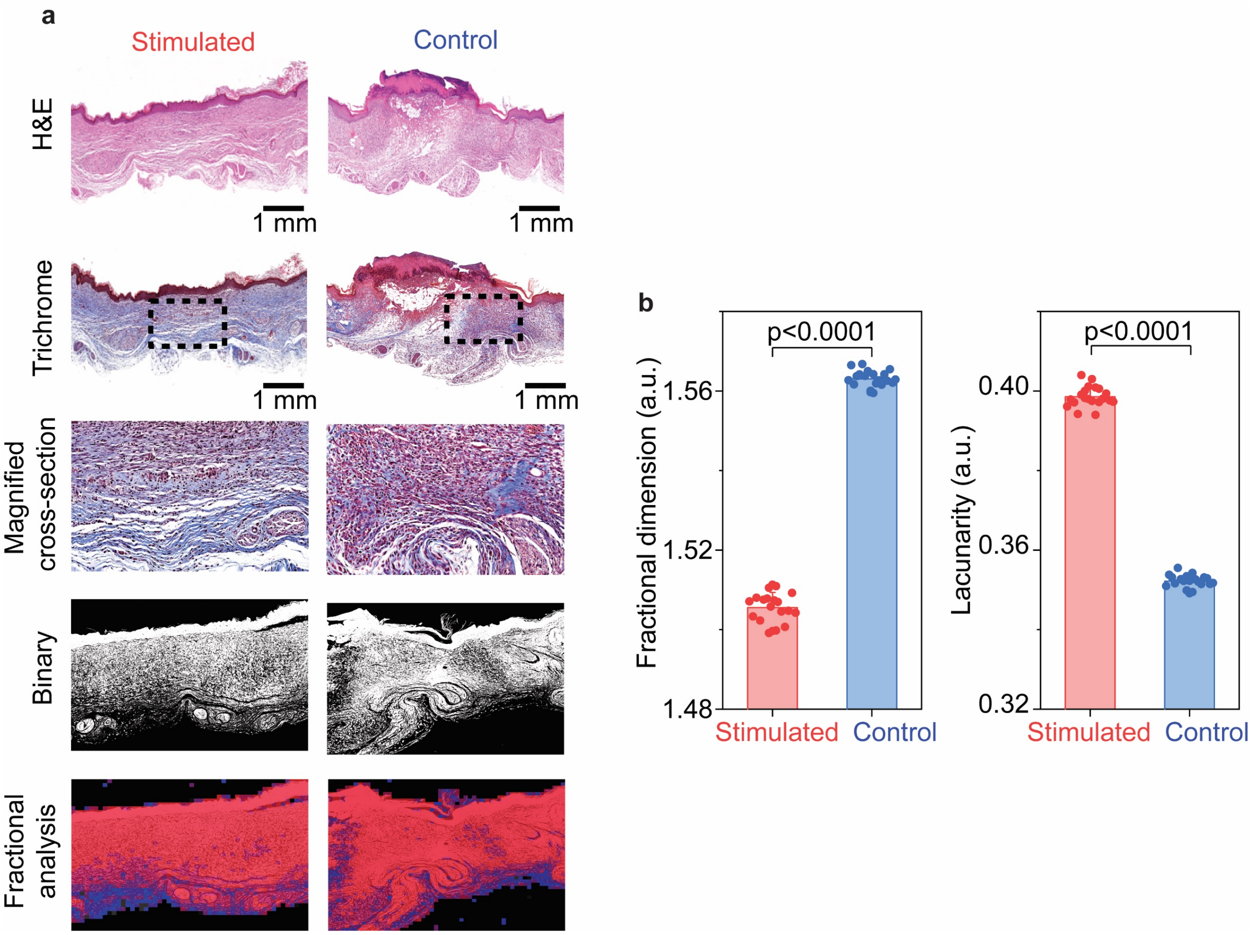

**Figure S27 | FracLac evaluation shows less scar-like fiber arrangement for STZ-induced diabetic wounds treated with electrical stimulation after 15 days. a,** Representative cross-sectional histology images and corresponding FracLac of skin tissues harvested from mice with and without electrical stimulation. Magnified cross-section, outlined with a dotted rectangle, highlights the collagen composition and microvessel formation of healed tissue. Masson's trichrome images were binarized using FracLac and collagen denisty was quantified, represented by red and blue colored boxes in the bottom-most row. **b,** Quantitative comparison of local fractal dimension (FD) and lacunarity. FD measures density of collagen networks. A higher FD has a denser and scar-like fiber arrangement. Lacunarity measures the amount of randomness or heterogeneity in a sample. A low lacunarity implies less heterogenous collagen fiber orientation. All data are represented as mean ± standard deviation. Two-tailed t-test assuming equal variances were performed for the *p* values.

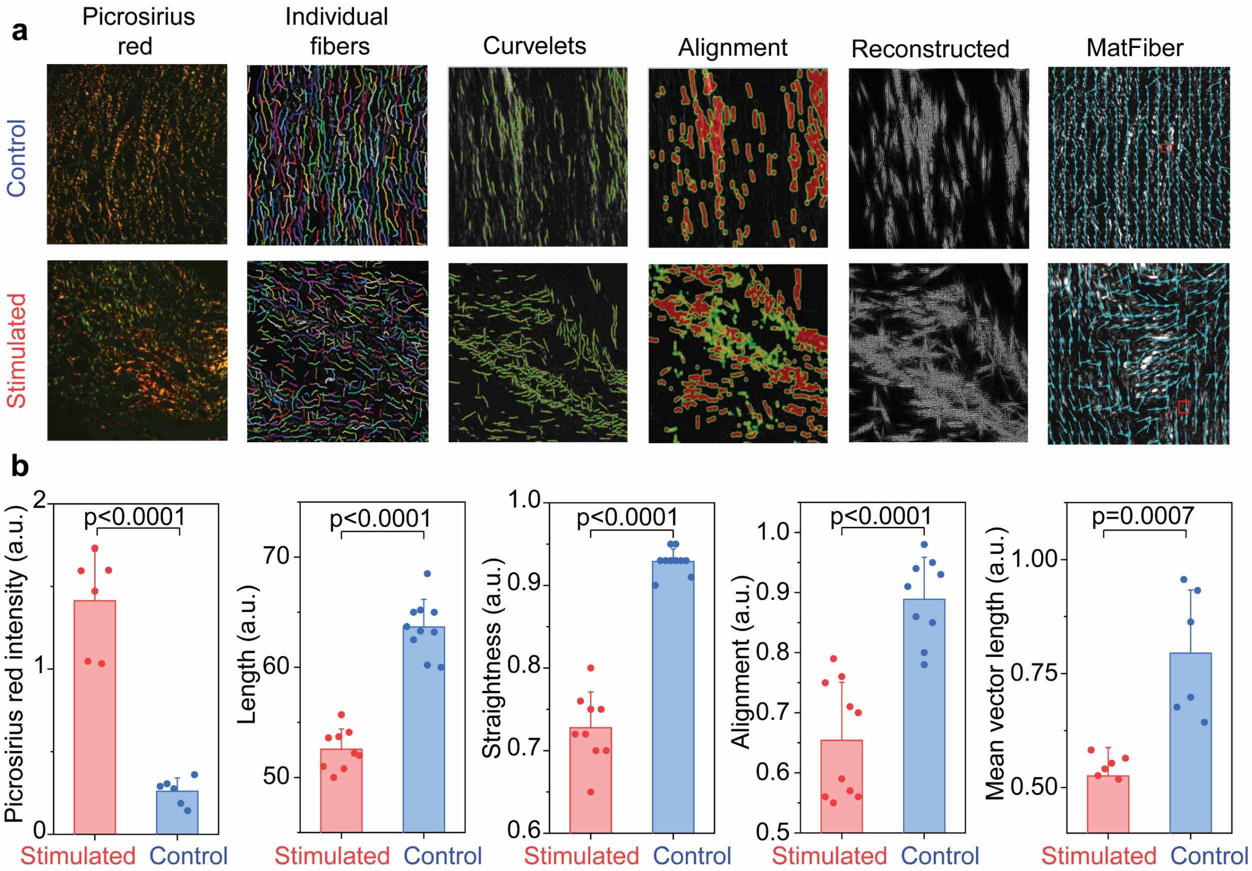

**Figure S28 | Picrosirius red staining shows more random collagen networks for STZ-induced diabetic excisional wounds treated with electrical stimulation after 15 days. a,** Representative picrosirius red stained images and collagen network analysis of skin tissues harvested from mice with and without electrical stimulation using Curve Align, CT-Fire and MatFiber. CT-FIRE algorithm outlines individual fibers to evaluate length, width, angle, and curvature. CurveAlign quantifies all fiber angles and strength of alignment within an image, tracing the fibers in green quantifying the fiber density with red heat maps and reconstructing the image in binary form to produce an output. MatFiber quantifies fiber alignment, tracing the fibers with blue arrows, reported as mean vector length. The strength of alignment ranges from a value of 0 (completely random fiber alignment) to 1 (completely aligned fibers). **b,** Quantitative comparison of picrosirius red intensity, collagen length, collagen straightness, collagen alignment, and mean vector length shows more random collagen networks and less fibrotic tissue formation. All data are represented as mean ± standard deviation. Two-tailed t-test assuming equal variances were performed for the *p* values.

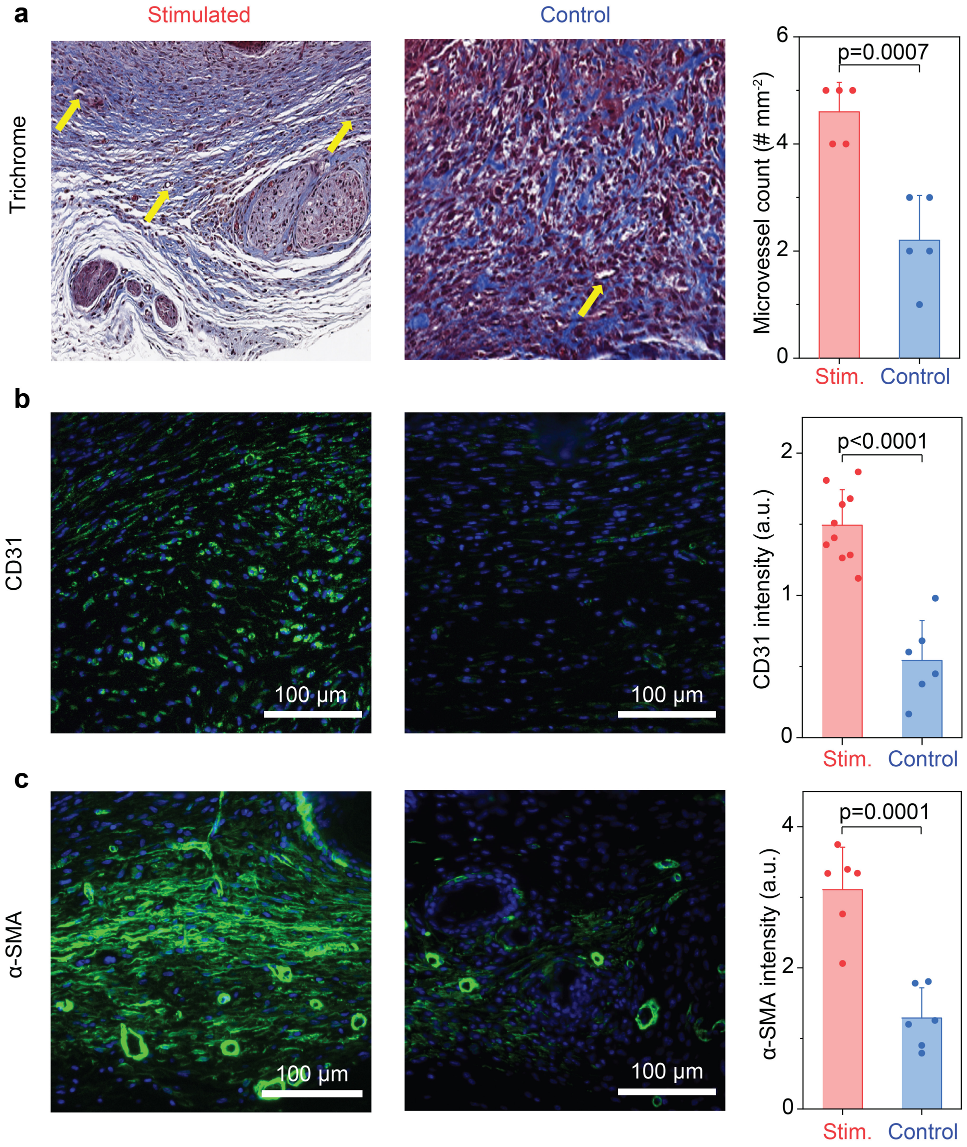

**Figure S29 | Histology and immunostaining reveals more neovascularization for STZ-induced diabetic excisional wounds treated with electrical stimulation after 15 days.** Representative Masson's trichrome (**a**), stained images and corresponding analysis showing microvessel count (indicated with yellow arrows), and vascularity of skin tissues harvested from mice with and without electrical stimulation stained with (**b**), CD31, which stains for blood vessels and (**c**), α-SMA, a marker for smooth muscle cells, namely myofibroblasts, which are commonly found in the walls of blood vessels. All data are represented as mean ± standard deviation. Two-tailed t-test assuming equal variances were performed for the *p* values.

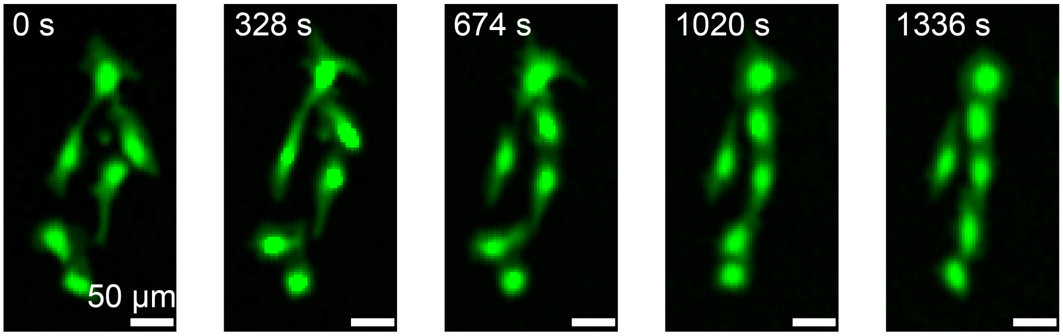

**Figure S30 | Fluorescent imaging showing human umbilical vein endothelial cells aligning under electric field (1 V cm$^{-1}$).**

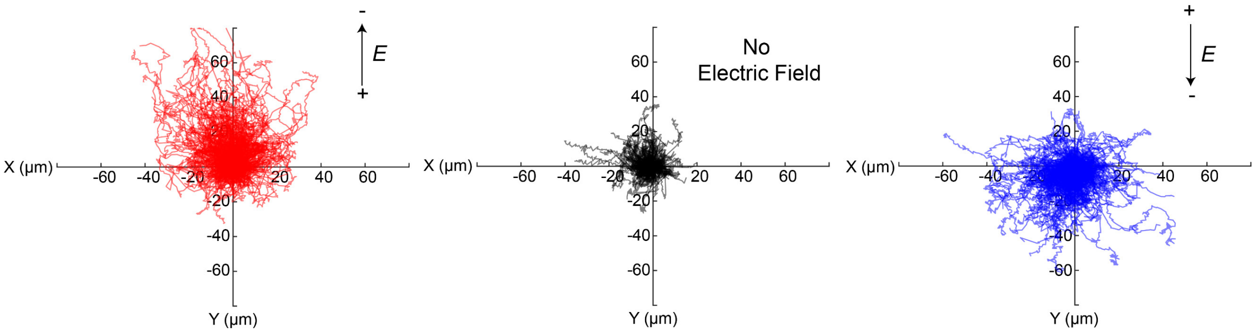

**Figure S31 | Trajectories of individual human umbilical vein endothelial cells showing directional migration under electric field (1 V cm$^{-1}$).** Cell trajectories were collected for 2 hours for each condition. Cells were maintained in basal media for the experiement.

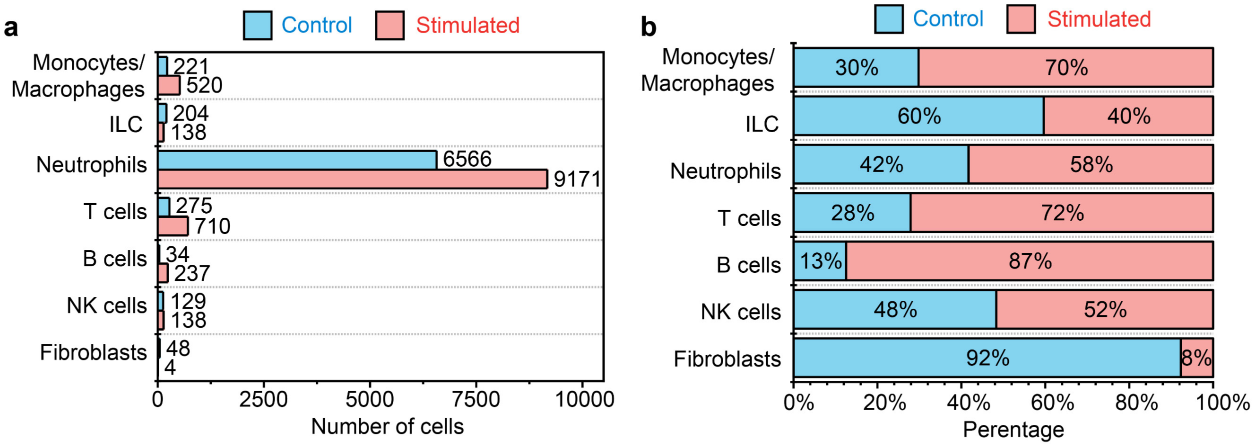

**Figure S32 | Quantitative comparison of the total number of cells from tissue with and without stimulation after 5 days (a) and relative percentages for each cell type (b).**

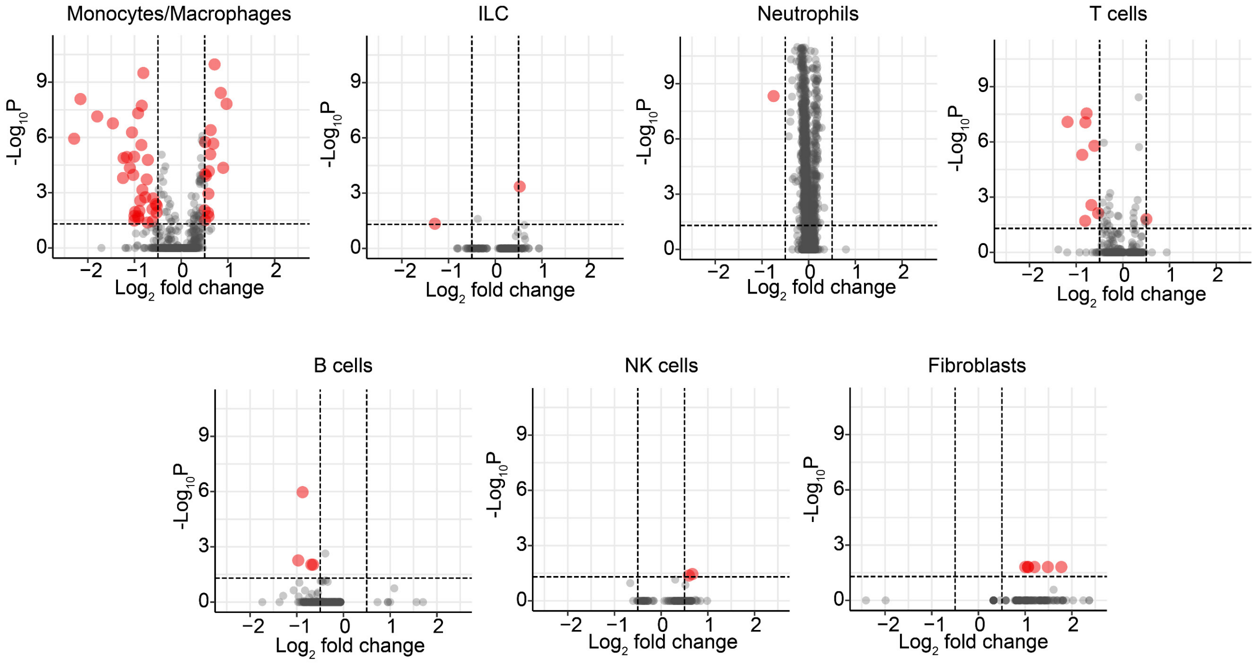

**Figure S33 | Volcano plots showing the number of genes with significant differences for each cell type.** Grey dots represent individual genes either above a certain log fold change or *p* value, while red dots represent genes both significantly different and with a high log fold change between stimulated and untreated wounds. Genes to the right of the *y* axis represent genes differentially upregulated to stimulated wounds, while genes to the left of the *y* axis represent differentially upregulated genes in untreated wounds. Macrophages/monocytes demonstrated the highest number of differentially upregulated genes in the stimulated group. *p* value cut off is 0.05 and fold change cut off is 0.5 for all groups. An overlaid volcano plot combining all cell types with color coded dots is presented in **Fig. 5c**.

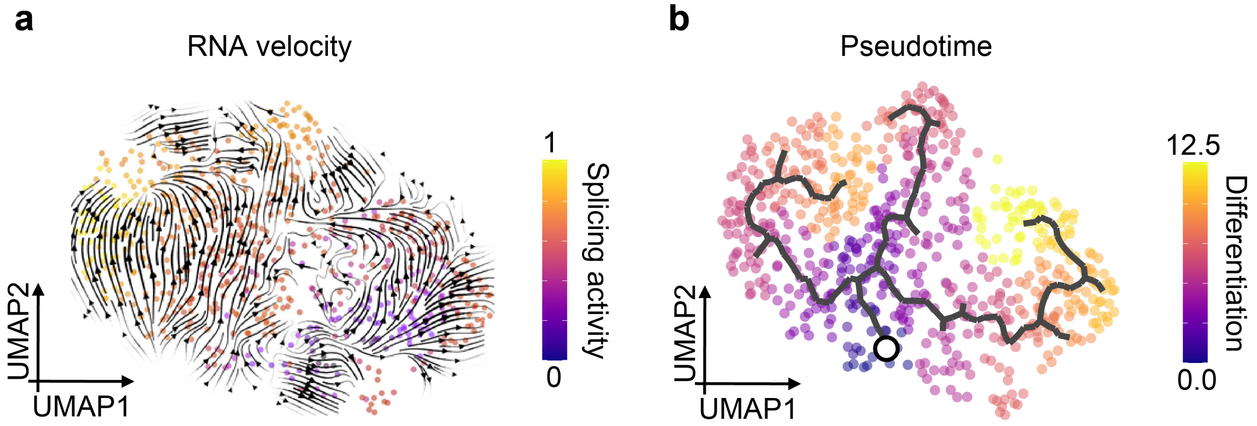

**Figure S34 | Pseudotime plot showing the trajectory of RNA velocity. a,** RNA velocity stream plot colored by latent time as computed by scVelo across all genes, quantifying overall differences in transcriptional dynamics[18]. Three general vector paths are identified for mRNA cell splicing activity with a relatively higher amount of differentiated individual cells found on the leftmost part of the plot (higher in latent time, more yellow-like hue and greater levels of mature spliced mRNA), and less differentiated cells found on the right (lower in latent time). **b,** Using Monocle 3, we identified an initial root node of differentiation (marked by a white circle) corresponding to our scVelo RNA velocity analysis that led to three trajectories of differentiation in the macrophage and monocyte subset shown in **Fig. 5e**. Cells at the initial state have a lower pseudotime value (denoted with a purple hue). As the distance from the initial state chronologically progresses towards the terminal state, the color transitions to a more yellow hue.

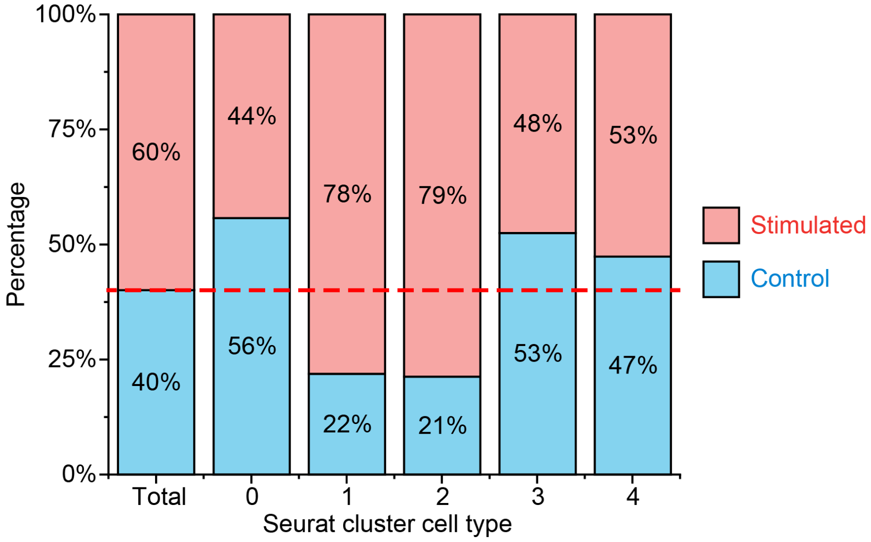

**Figure S35 | Relative percentage showing the ratio of the number of cells in the stimulated and control groups for each Seurat cluster in the macrophage and monocyte group.** For clusters 1 and 2, the stimulated group was predominant, as compared to control. The Seurat cluster is presented in **Fig. 5f.**

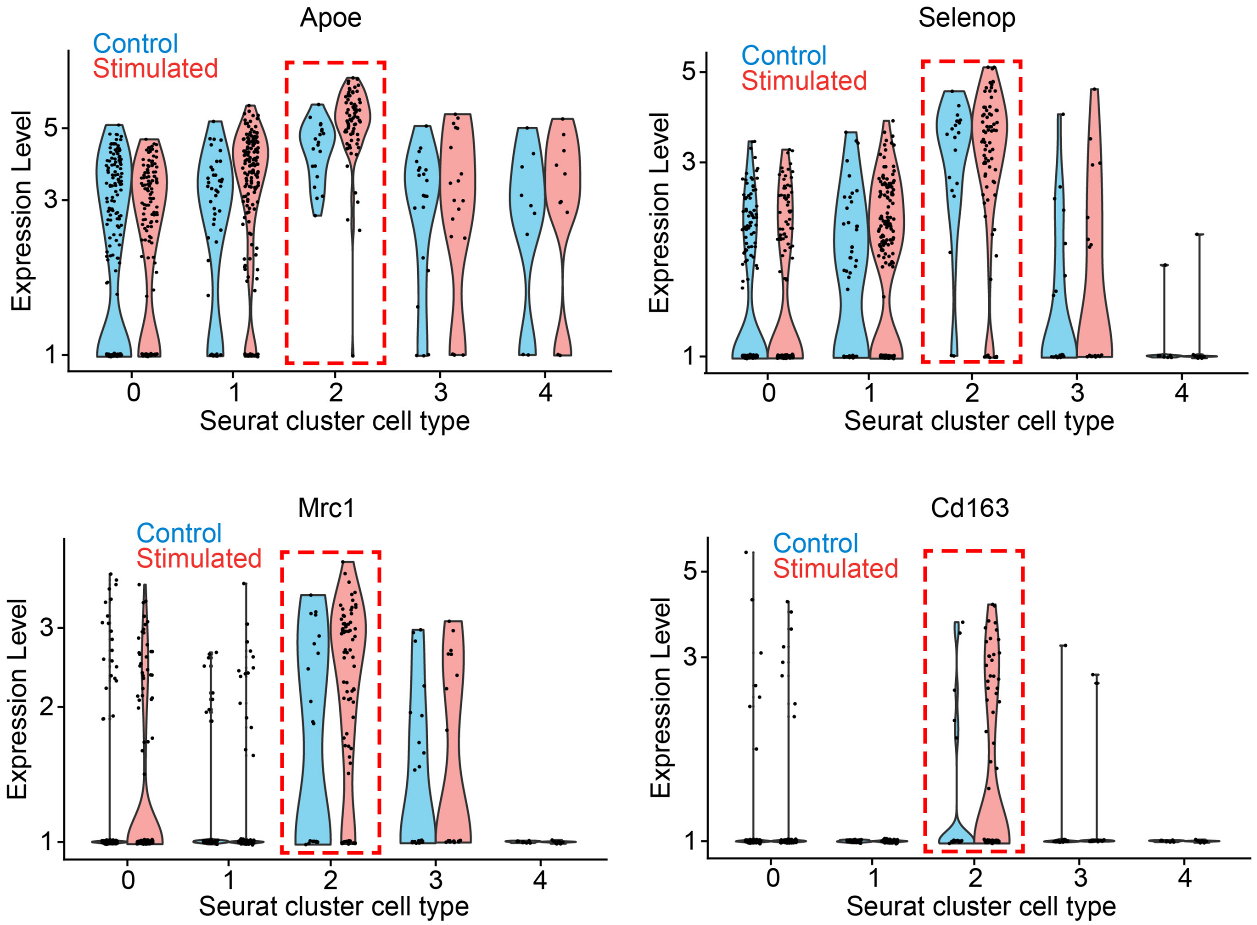

**Figure S36 | Violin plots showing the relative expression profiles of pro-regenerative genes in the stimulated and control groups within the macrophage and monocyte Seurat cluster.** An increased expression level was observed predominanlty in cluster 2, associated with predominantly pro-regenerative macrophage and monocyte subpopulations. Feature plots are presented in **Fig. 5h.**

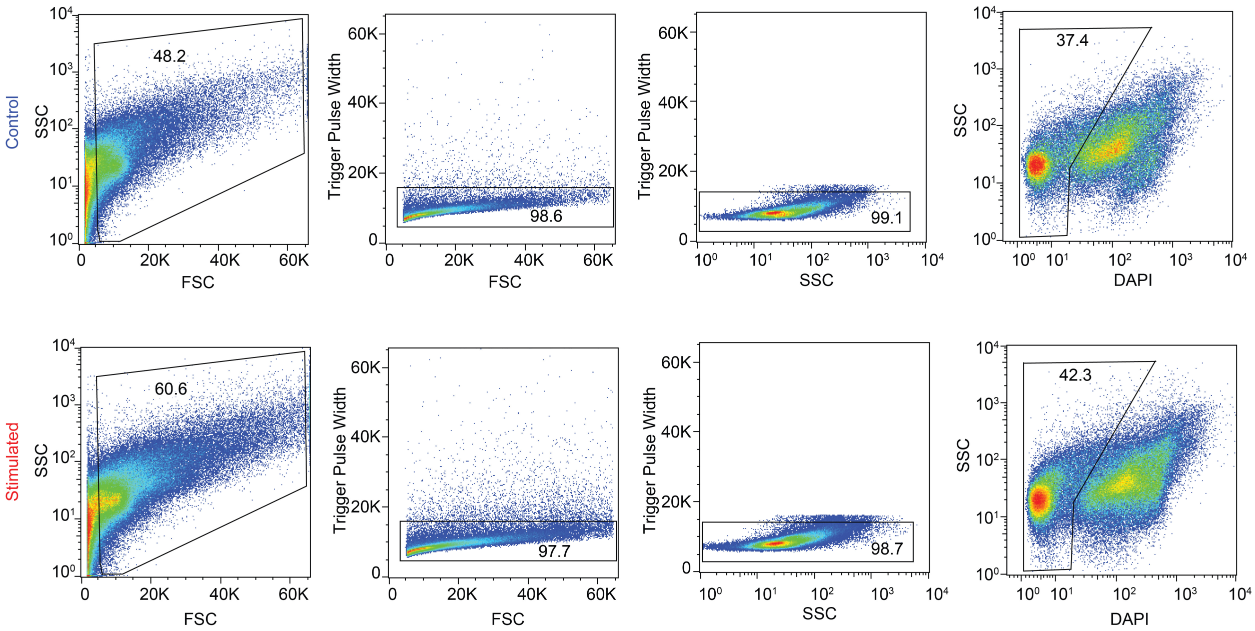

**Figure S37 | Gating strategy for the fluorescence-activated cell sorting (FACS) of cells in the excisional wound of the wild-type mouse in the GFP$^+$/WT parabiosis model.** GFP$^+$ cells were further gated as shown in **Fig. 5i.** SSC: side scattering, FSC: forward scattering, DAPI: 4',6-diamidino-2-phenylindole.
